# Treating Myotonic Dystrophy with artificial RNA endonucleases to specifically degrade toxic RNA expansions

**DOI:** 10.1101/2025.06.10.658795

**Authors:** Tong Wei, Hanyang Hu, Xin Jiang, Yucheng Li, Wenjian Han, Yi Yang, Xiao Xiao, Miaowei Mao, Zefeng Wang

**Affiliations:** School of Life Science, Guangming Advanced Research Institute, Southern University of Science and Technology, Shenzhen, Guangdong, China; Shanghai Institute of Immunity and Infection, Chinese Academy of Sciences, Shanghai, China; Shanghai Institute of Nutrition and Health, Chinese Academy of Sciences, Shanghai, China; Belief BioMed, Shanghai, China; School of Pharmacy, East China University of Science and Technology, Shanghai, China; Songjiang Research Institute, Songjiang Hospital, Shanghai Jiao Tong University School of Medicine, Shanghai, China

## Abstract

Myotonic dystrophy type 1 (DM1), the most common autosomal dominant muscular disorder, is driven by expanded CUG repeats in the 3′ UTR of DMPK gene, which sequester RNA-binding proteins and cause aberrant alternative splicing. Despite extensive study, no effective treatment exists, and current care remains limited to symptomatic management. Using a customized dual-color reporter carrying RNA repeats in different lengths, we engineered a new class of programmable artificial enzymes (artificial RNA cleavers, or ARCs) that preferentially degrade the pathogenic repeats both in cells and in a mouse model. We further examined the *in vivo* efficacy of ARCs by systematically delivering ARCs into the DM1 mouse model with muscle-specific adeno-associated viruses, and validated that ARCs can rescue the muscular pathology and movement ability. The treatment by ARCs showed sustained efficacy, low immunogenicity, and minimal off-target effects in animal model, which are key concerns in clinical translation of gene therapies. Collectively, these findings establish ARCs as a potent and precise RNA-targeting platform with strong clinical potential for the treatment of DM1 and other repeat expansion disorders.

## INTRODUCTION

The advances of sequencing technology have led to the discovery of an increasing number of genetic diseases ^1,2^, among which a major class of genetic disorders is caused by the abnormal expansion of short tandem repeats ^2^. The variable lengths of these repeat expansions enable them to act as heritable pathogenic mutations *via* multiple mechanisms, including epigenetic silencing, RNA metabolism disruption, and gain-of-function mutations in proteins ^3^. More than 50 of such autosomal dominant disorders have been identified to date, including myotonic dystrophy (DM1 and DM2), Huntington’s disease (HD), and C9orf72-associated familial amyotrophic lateral sclerosis (ALS) ^4–7^.

DM1 is the most common form of myotonic dystrophy, with recent surveys indicating an estimated prevalence up to ∼1/2,000 in population ^8,9^, much higher than what reported previously ^10^. This disease is caused by an abnormal (CTG)_n_ tandem repeat (n > 50) in the 3’ UTR of *DMPK* gene ^9^. Transcripts from this pathogenic locus contain long <u>C</u>UG repeat <u>e</u>xpansions (CREs) that sequester RNA-binding proteins (RBPs) such as MBNL1 and CUGBP1, leading to RNA-protein aggregates in nucleus ^11–13^. These aggregates could severely disrupt normal RNA metabolism of muscle cells, leading to myotonia, muscle degeneration, cardiac conduction defects, and respiratory insufficiency, which can significantly affect the mortality rate of patients ^14,15^.

Despite extensive insights into DM1 pathogenesis, the FDA only approved limited therapeutic interventions, which primarily alleviate symptoms rather than target the underlying cause — CREs. Several new strategies have been developed to target CREs at the DNA or RNA level. At DNA level, the ideal approach is to remove the pathogenic expansions using genome editing tools, which has the potential to cure such diseases ^16,17^. However, due to the unique feature of the expansion region, this method actually causes DNA breaks and may lead to expansion growth during the DNA repair process, exacerbating the progression of the disease ^17–19^. In addition, genome editing tools are generally unable to distinguish between normal and pathogenic repeat regions, and thus likely to affect the functional activity of normal alleles. In the past decade, manipulating the CREs at the RNA level has also become a promising strategy for gene therapy, such as blocking the transcription of pathogenic regions ^20^, disrupting interactions between the CREs and RBPs ^21–23^, or degrading pathogenic RNA directly ^24–28^. These approaches include modified antisense oligonucleotides (ASOs) ^28^, RNA interference with shRNA ^29^, antibody-siRNA conjugates ^30^, antibody-ASO conjugates ^31,32^, dSaCas9-mediated transcriptional inhibition ^20^, small molecules-^22^ or RBPs ^21,23^-mediated competitive binding of CREs, CREs degradation mediated by chemical compounds ^33^ or therapeutic proteins (e.g., PUF-PIN ^25^; PIN-RCas9 ^26,34^ or Cas13d ^27^) (Supplementary Table 1).

However, lack of head-to-head comparations of these approaches makes it unclear to assess their efficacy. Moreover, each of these methods has its own limitations, such as inefficient delivery for ASOs ^35^, low targeting specificity for small molecules ^22^, and high immunogenicity of foreign proteins like Cas proteins ^36^. Two state-of-the-art gene therapy approaches, PIN-RCas9 ^34^and Cas13d ^27,37^, face practical concerns for clinical translation. The PIN-RCas9 system has a large size that complicates delivery (two viral vectors have to be used together ^34^); whereas the Cas13d exhibits excessive RNase activity that produce broad off-target effect including degradation of wide-type short RNA repeats ^38^. In addition, the CRISPR/Cas-based approaches using bacterial proteins also have to deal with the pre-existing immunity in humans ^36,39,40^, thus requiring simultaneous immunosuppressive treatment ^34,41^. The artificial RNA endonucleases previously developed by our group using human proteins showed low efficacy in animal models ^42^. Therefore, it is highly desirable to develop efficient and practical treatments for CRE disorders that are life-threatening to thousands of patients.

In this study, with a dual-color reporter expressing varying lengths of CREs, we engineered and optimized a series of RNA endonucleases called artificial RNA cleavers (ARCs) that specifically and efficiently degrade CREs. Additionally, we evaluated the available therapeutic strategies targeting CRE regions using cultured cells and an animal model. Our results demonstrate that ARCs outperform other methods in efficiently degrading toxic CREs and reducing nuclear RNA foci, subsequently releasing MBNL1 to restore its normal cellular localization. The *in vivo* efficacy of ARCs was further validated in a well-established DM1 mouse model. The ARC system has multiple advantages, including length preference for pathogenic RNAs, programmability, low immunogenicity, suitable size for delivery by tissue specific-AAVs (adeno-associated viruses), highlighting its potential as a promising therapeutic strategy in treating RNA diseases caused by CREs.

## RESULTS

### Design and optimization of the ARCs

To mimic the genetic features of DM1, we developed a reporter system containing different numbers of CUG repeats. This reporter was designed to express two distinct mRNAs: the EGFP mRNA containing a pathogenic CRE in its 3’ UTR, and the mCherry mRNA containing a non-pathogenic short (CUG)_12_ repeat as the control (Fig. 1a). Inspired by the design of ASREs ^42^, we fused an engineered PUF domain (RNA binding scaffold of Pumilio and FBF homology) recognizing CUG repeats ^25^ with different RNA endonucleases to screen a series of artificial enzymes (artificial RNA cleavers, noted as ARCs). To assess the degradation efficiency of CREs, we co-transfected the ARCs with the dual-color reporter in HEK 293T cells. Compared to the control, we found that several fusion proteins showed efficient degradation of pathogenic RNAs while having much smaller impact on normal RNAs (Fig. 1b). Due to the therapeutic requirement (e.g., excessive activity could impact the transcriptome, while large protein size may pose challenges for delivery), we chose the PUF-SUPV3L1 and RNASE T2-PUF as candidates for subsequent optimization, as both enzymes showed a length-dependent degradation of CREs.

**Fig. 1.**
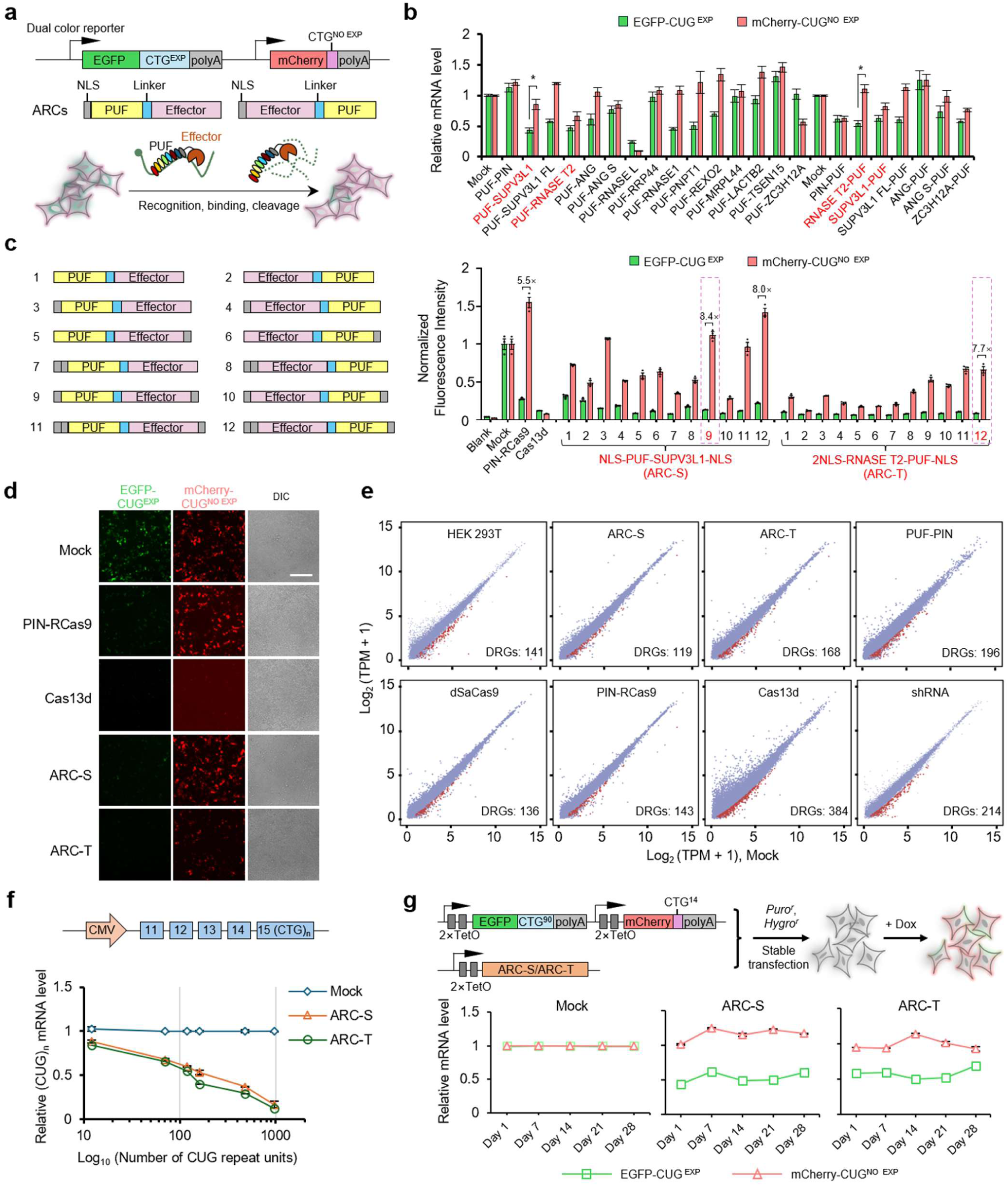
Engineered artificial enzyme to degrade pathogenic CREs. **a**, Overview of ARCs design and characterization. Top, the dual-fluorescence construct expressing CRE and non-CRE RNAs in the 3’ UTR of EGFP and mCherry respectively. Middle, domain configuration of ARCs. NLS, nuclear localization signal. Bottom, schematic of ARCs-mediated specific RNA degradation within the dual-fluorescence system. **b,** Relative mRNA abundance in HEK 293T cells co-transfected with dual-fluorescence reporters and various ARC-encoding plasmids. EGFP and mCherry RNA levels were normalized to the vector control (Mock). Data are mean ± s.e.m. (n=3). **c,** Schematics of ARCs optimization strategy (left). Relative fluorescence intensities of the ARCs compared to Mock group were plotted, with the fold change indicated on top. Data are mean ± s.e.m. (n=3). **d,** Fluorescence images of dual fluorescence changes in co-transfected HEK 293T cells for different methods. Scale bar, 200 μm. **e,** Scatter plots for gene expression levels between Mock and different treatment groups. The mock group represents cells transfected only with the reporter vector and the control vector. The down-regulated genes (DRGs, adjusted *P*< 0.05, log_2_FC < −1) were marked in red. TPM, transcripts per million. **f,** The *DMPK* minigene reporters containing varying lengths of (CUG)_n_ were transfected into HEK 293T cells after ARCs transfection, and the relative mRNA level (normalized to the Mock transfection) were plotted. Statistical significance was assessed by an unpaired two-tailed Student’s *t*-test (\**P*< 0.05). **g,** Diagram illustrating a tetracycline-inducible system for dual-color based assay to assess the long-term activity of ARCs (top). The dual-fluorescence reporter and ARCs constructs were co-transfected, and the cells were selected using Hygromycin (*Hygro*^r^) and Puromycin (*Puro*^r^) resistance, respectively. Quantification of the CRE and non-CRE RNA abundance in *Hygro*^r^ and *Puro*^r^ positive cells (bottom). TetO, tetracycline operator. Dox, Doxycycline.

We further improved the performance of two ARC candidates by screening inter-domain linkers and rearranging the domain configuration. We tested various peptides from the Linker database ^43^ by inserting these linkers between the PUF domain and functional modules, and identified Linker 3 as the optimal linker for further optimization (Extended Data Fig. 1a and 1b). The number of NLS (nuclear localization signal) and the configuration of fusion proteins were also investigated, leading to varying improvement in the CREs degradation by ARCs as judged by the dual-fluorescence reporter (Fig. 1c). In particular, NLS-PUF-SUPV3L1-NLS (noted as ARC-S) and 2NLS-RNASE T2-PUF-NLS (noted as ARC-T) exhibited an approximate ∼8-fold difference in relative changes of fluorescent intensity with PIN-RCas9 ^26^ and Cas13d ^27^ as the positive controls (Fig. 1c). We also used fluorescence microscopy to compare ARCs with the PIN-RCas9 and Cas13d, and found that PIN-RCas9, ARC-S and ARC-T exhibited improved length preferences on CREs, whereas Cas13d simultaneously eliminated both red and green fluorescence signals (Fig. 1d). The activities of ARC-S and ARC-T were further validated in a cell line stably expressing tetracycline-inducible reporters, producing consistent results (Extended Data Fig. 1c).

Since multiple methods have previously been reported to degrade the CREs of DM1, we conducted a parallel comparison to comprehensively evaluate the efficiency and specificity of these tools (Fig. 1e and Extended Data Fig. 2). We confirmed that ARC-S, ARC-T and PIN-Rcas9 can preferably degrade long CREs, while the Cas13d and shRNA degraded both short and long CUG repeats in a similar efficiency (Extended Data Fig. 2b). In addition, the RNA-seq analysis revealed the PIN-RCas9, ARC-S and ARC-T have less off-target effect on transcriptome compared to the other existing tools (Fig. 1e and Extended Data Fig. 2). Collectively, these results showed that the ARCs system outperformed the other methods as a potential gene therapy strategy in the cultured cells.

To further assess the degradation capability of ARCs for CREs of varying lengths, we co-transfected ARCs with plasmids expressing various numbers of CUG repeats (0, 12, 70, 160, 480, and 960) into HEK 293T cells and quantitatively measured the residual CRE levels. As hypothesized, we observed a length dependent degradation of (CUG)_n_ repeats for the ARC system (Fig. 1f). The cleavage efficiency of the ARC-S and ARC-T improved with the increased length of CREs, possibly due to the enhanced recruitment of ARCs to the target RNAs. Additionally, we found that ARC-S and ARC-T can achieve long-term reduction of pathogenic RNA level in a tetracycline-inducible cell line (Fig. 1g), indicating their therapeutic potential for RNA diseases caused by CREs.

### Elimination of pathogenic RNA foci in living cells

The pathogenic RNA expansion usually led to the nuclear aggregation of RNA/RBP as nuclear foci, which are generally visualized using RNA FISH (fluorescence *in situ* hybridization) in fixed cells. However, the numbers of nuclear foci detected by RNA FISH are often affected by different experimental conditions (such as the wash procedure and fluorescence probes). Compared to FISH, the RNA imaging in living cells is more precise and reliable, especially for better quantification of RNAs ^44^. To enable monitoring RNA targets in living cells, we labelled CREs by inserting fluorescent RNA aptamer (Pepper) to DMPK minigene reporter ^45^, which has different numbers of CUG repeat units (12 and 960 units) at the 3’ terminal. Repetitive RNAs can be detected by adding a non-fluorescent dye, which becomes fluorescent once it is locked within Pepper aptamer ^45^ (Fig. 2a).

**Fig. 2.**
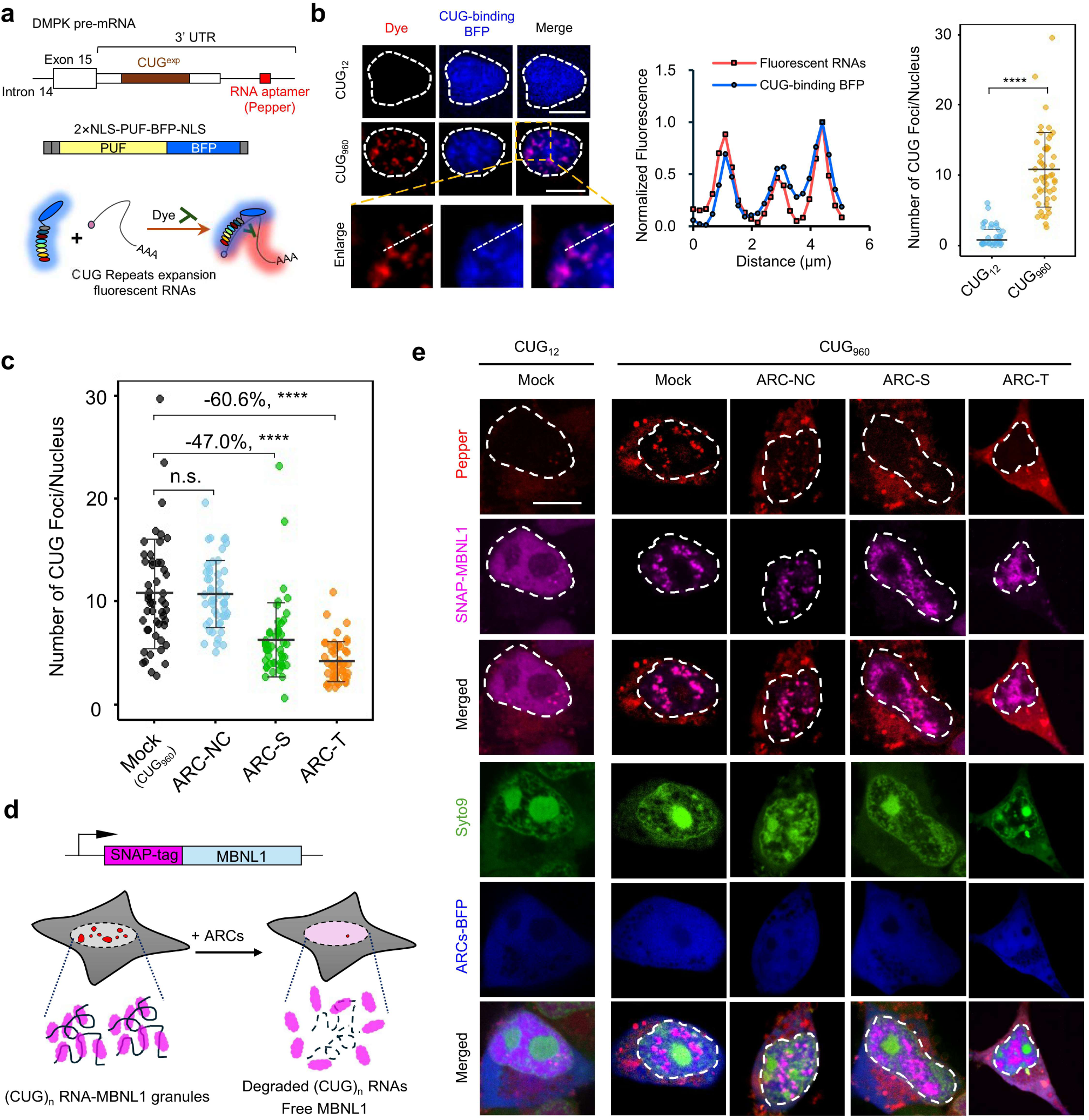
ARCs reduce RNA foci and restore MBNL1 localization in living cells. **a**, Schematic of the *DMPK* pre-mRNA containing CTG repeat tracts and RNA aptamer (Pepper) recognized by the PUF-BFP fusion protein. **b,** Live-cell imaging of HEK 293T cells expressing the PUF-BFP fusion protein and fluorescent Pepper containing 12 or 960 CTG repeat tracts (left). The nuclear regions were indicated with dashed lines. Scale bar, 5 μm. Middle, fluorescence intensity profiles of Pepper and BFP signals along the indicated dotted line. Right, statistical analysis of the numbers of RNA foci in cells expressing 12 or 960 CTG repeat tracts. **c,** The numbers of RNA foci in different groups. The degradation efficiencies of ARC-S and ARC-T were labeled above the lines. The mean ± s.d. from 50 representative cells were plotted. **d,** Schematic of ARCs-mediated degradation of CRE RNAs and release of sequestered SNAP-MBNL1. **e,** Live-cell imaging of RNA foci (red) and MBNL1 (magenta) localization in the ARCs treated cells stably expressing SNAP-MBNL1. The nuclear regions were stained with Syto9 (green). Scale bar, 5 μm. Statistical significance was assessed by an unpaired two-tailed Student’s *t*-test (\*\*\*\**P*< 0.0001; n.s., not significant).

We simultaneously monitored nuclear-localized PUF-BFP fusion protein that was programmed to recognize CUG repeats, and found that PUF-BFP exhibit identical positions with the CRE-Peppers (Fig. 2b). Consistent with previous studies ^46,47^, the cells with CUG_12_ RNAs have essentially no accumulated foci (<1 foci/cell), whereas CUG_960_ RNAs can be detected with ∼10 RNA foci in each nucleus (Fig. 2b). Using this strategy, we systematically evaluated the capacity of existing tools to remove RNA foci in living cells, and found that most methods (except PUF-PIN) significantly reduced toxic RNA foci, with the ARC-T demonstrated the highest efficiency in clearing RNA foci (Fig. 2c and Extended Data Fig. 3).

Next we investigated the subcellular localization of endogenous CUG-binding proteins by generating a cell line stably expressing SNAP-tagged MBNL1 ^48^ (Fig. 2d). Previous reports showed that MBNL1 can be sequestered by CREs in the nuclear foci, leading to a loss of its normal function in regulating RNA splicing ^11,21,33^. Using the live-cell imaging, we found that MBNL1 exhibited normal nuclear distribution in the presence of CUG_12_, whereas it was sequestered by pathogenic RNA foci in the presence of CUG_960_ (Fig. 2e). This is the first time to visualize MBNL1 and CUG repeats of varying lengths in living cells. Notably, the expression of ARC-S or ARC-T can mitigate the nuclear aggregation of MBNL1 while eliminating the RNA foci (Fig. 2e), highlighting the ability of ARCs to reduce RNA foci and restore normal localization of MBNL1.

### Degradation of toxic RNA repeats by ARCs in a new mouse model

We further evaluated the ability of ARCs to degrade pathogenic CRE RNAs with a new animal model that allows quick assessment of CREs cleavage *in vivo*. To this end, we generated a <u>D</u>ual <u>C</u>olor <u>C</u>UG (DCC) transgenic mouse model from C57BL/6 strain, where a similar dual-fluorescence reporter containing different (CUG)_n_ repeats was integrated into the *Rosa26* locus (Fig. 3a and Extended Data Fig. 4). The global difference of gene expression between DCC and wild-type mice was first determined using RNA-seq. In total, we found approximately 300 genes with altered expressions (122 genes upregulated and 174 genes downregulated in DCC mouse) or alternative splicing (AS) events (335 AS events, FDR < 0.05, |Δ PSI|> 0.2) in the quadriceps (Extended Data Fig. 5). As expected, these altered genes were functionally enriched in pathways related to the development of the muscular system and metal ion transport (Extended Data Fig. 5). However, such changes on the transcriptomic level only slightly altered certain behaviors or motor ability in DCC mice (Extended Data Fig. 6). Nevertheless, this model still represents an *in vivo* platform for rapid evaluation and comparison of the potential CRE-targeting approaches.

**Fig. 3.**
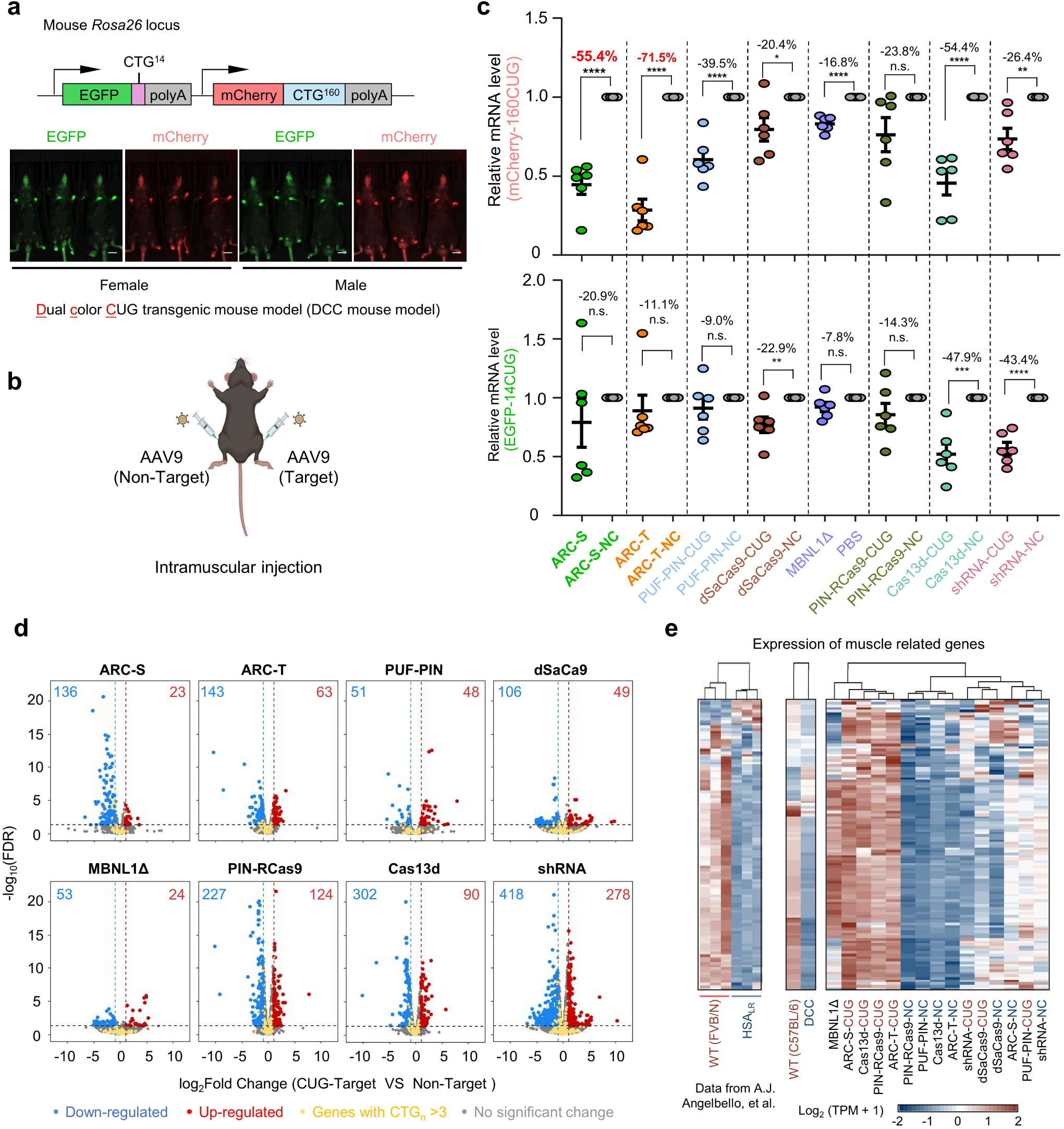
ARCs exhibit therapeutic efficiency in DCC mice. **a**, Schematic of the <u>D</u>ual <u>C</u>olor <u>C</u>UG^exp^ (DCC) transgenic mouse model (top) and the fluorescence imaging of the mice (bottom). Scale bar, 1 cm. **b,** Treatment scheme for AAV9-mediated delivery of current therapeutic tools. A pair of vectors encoding different CRE-targeting tools or the corresponding non-targeting control were injected into the right and left quadriceps of DCC mice (aged 6-8 weeks). **c,** Levels of CRE and non-CRE transcripts in DCC mouse muscle were quantified by RT-qPCR at 8 weeks post intramuscular injection. Relative mRNA level was normalized to the corresponding non-targeting control. The numbers above the lines represented degradation efficiency. Data are mean ± s.d.(n=6). **d,** Global analysis of gene expression in AAV-treated quadriceps of DCC mice. Yellow dots indicate genes with ≥3 CUG repeats, while red and blue dots represent up- and down-regulated genes (adjusted *P* < 0.05, |log_2_FC| > 1). **e,** Hierarchical clustered heatmap shows normalized gene expression levels of muscle-related genes. Left, identification of the differentially expressed genes (DEGs) in skeletal muscle from HSA_LR_ *vs.* WT mice. Right, expression patterns of the same DEGs across WT mice, DCC mice and experimentally treated DCC groups (n=3 for each group). Statistical significance was assessed by an unpaired two-tailed Student’s *t*-test (\**P*< 0.05; \*\**P*< 0.01; \*\*\**P*< 0.001;\*\*\*\**P*< 0.0001; n.s., not significant).

To systematically determine the *in vivo* efficacy of various therapeutic tools in targeting CREs, we used AAV2/9 vectors to deliver different genes using muscle injections. The PUF-PIN, ARC-S, ARC-T, MBNL1Δ (a truncated MBNL1 to bind CUG repeats as competitive inhibitor) ^21^, and a (CUG)_n_-targeting shRNA ^29^ were packed in a single AAV vector, whereas two separate AAV vectors were used to pack the CRISPR/Cas-based nucleases and their corresponding sgRNAs due to the limited capacity of AAVs ^34^. The AAVs were injected into the right quadriceps of 8-week-old DCC mice, with an equal dose of non-CUG targeting control AAVs injected into the left quadriceps (Fig. 3b). As expected, we found all treatments reduced the levels of mCherry-160CUG, with ARC-S and ARC-T exhibiting the highest activity (degrading 55% and 72% of the toxic RNAs) (Fig. 3c). Although the Cas13d also showed >50% degradation of mCherry-160CUG (∼54%) compared to the control, the ARCs had less impact on the control target (EGFP-14CUG) than the Cas13d (Fig. 3c), indicating that the ARCs are more effective in selectively cleave CREs. The high activity of Cas13d in degrading both long and short CUG repeats is consistent with a severe off-target effect on transcriptome as reported previously ^38,49^. In contrast, shRNA preferentially targeted short CUG sequences, indicating a weak ability to reduce nuclear CREs.

We further investigated the global off-target effects of these therapeutic tools using RNA-seq, and observed that the ARCs, PUF-PIN, dSaCas9 and MBNL1Δ exhibited smaller impact on gene expression (up to ∼200 genes affected), while PIN-RCas9, Cas13d and shRNA showed a larger effect on the transcriptomes (expression of 350-697 genes were affected, Fig. 3d). We further compared these data with the transcriptome of HSA_LR_ mice that were generated by systemically overexpressing ∼220 CTG repeats in muscles ^50^. The HSA_LR_ mice can recapitulate many of the muscle and mobility phenotypes in DM1 patients, and thus are widely used in therapeutic development. Using previously published RNA-seq data ^33^, we found that 113 muscle-related genes exhibited significant expression changes in the skeletal muscles between wild-type (FVB/N strain) and HSA_LR_ mice (Fig. 3e, left). Transcriptomic analyses confirmed that these muscle-related genes showed similar changes between the wild-type (C57BL/6 strain) and DCC mice, despite these two models being generated from different background strains (Fig. 3e, middle). More importantly, the treatment by CUG-targeting ARCs, PIN-RCas9 and Cas13d on DCC mice led to a general correction of the 113 muscle-related genes toward the expression pattern of the wild-type mice (Fig. 3e, right). As expected, the CUG-targeting shRNA and dSaCas9 did not show such pattern shift, consistent with the weak decrease of (CUG)_160_ by these methods (Fig. 3c). We also examined the change of AS events in DCC model treated by various therapeutic tools, and confirmed that the treatments by CUG-targeting ARCs had largely restored the splicing alteration in the DCC mice (Extended Data Fig. 7).

### Systematic delivery of ARCs with AAV vectors

The toxic RNA repeats in DM1 cause defects in multiple tissues with especially severe damage in muscles and heart, resulting in symptoms such as muscle weakness and myotonia ^51^. However, delivery of therapeutic agents to muscle tissues is challenging, and previously studies even combined local injections with *in vivo* electroporation to achieve efficient delivery of therapeutics ^52^, which is unpractical for patients. In addition, due to the pre-existing antibodies against bacterial proteins in human, the CRISPR/Cas-based approaches may induce severe immune responses in a large fraction of population ^36^.

To explore the therapeutic application of ARCs, we systemically delivered ARCs with AAV2/9 vector using tail vein injection in DCC mice. Considering potential immunogenicity, we first examined the immune response of animals to the ARC treatment at different time-points ^34,36^. After 6 and 12 weeks of AAV2/9-ARCs treatment, we collected the CD4^+^ and CD8^+^ T cells from fresh peripheral blood mononuclear cells (PBMCs) and measured the levels of immune markers (Fig. 4a). Our data suggested that the expression levels of the immune markers (IFN-γ, TNF-α, CD25, and CD107A) were essentially unchanged by ARC treatment, indicating that T cells were not stimulated by the treatment (Fig. 4b and 4c).

**Fig. 4.**
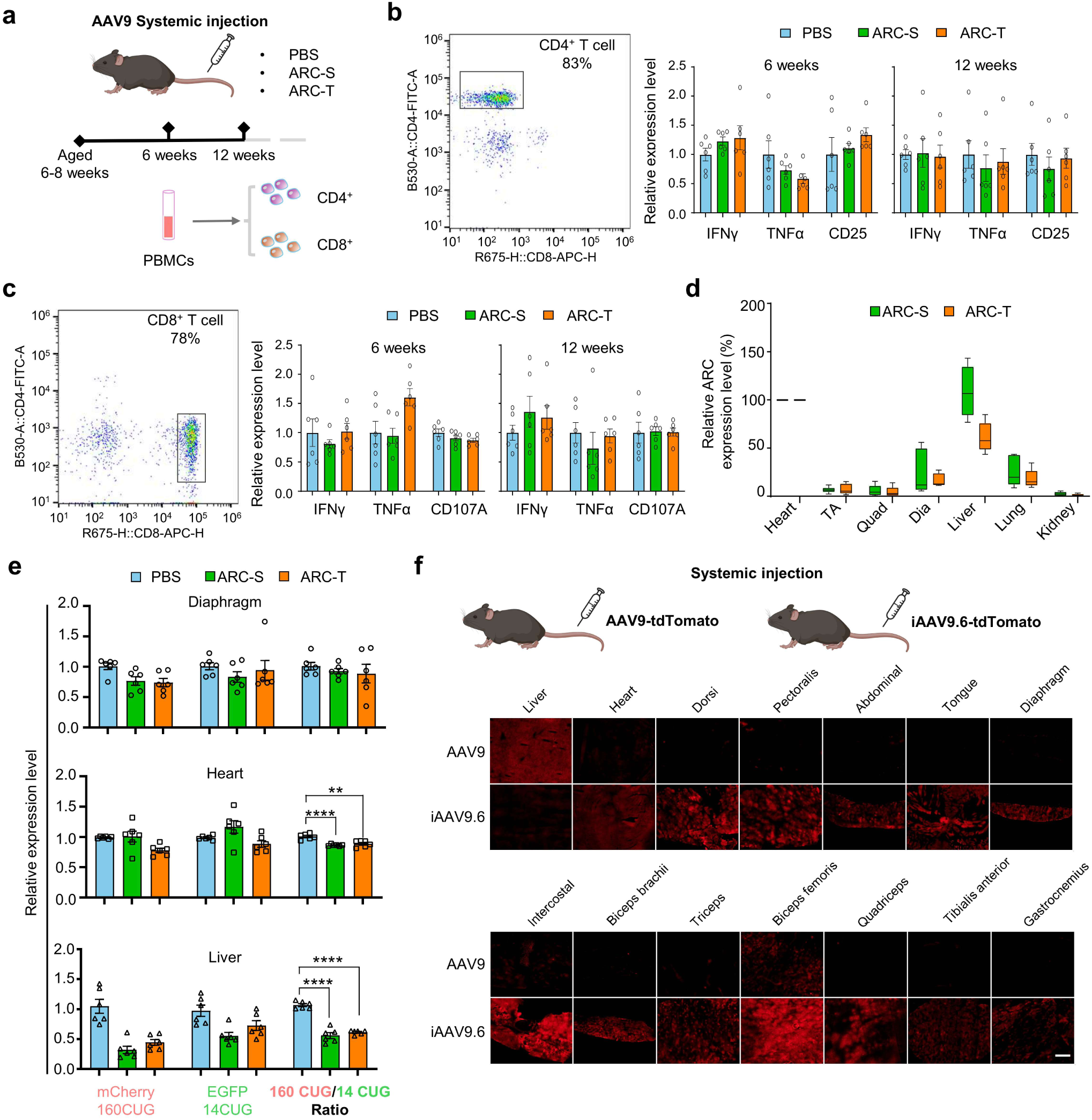
ARCs delivery and enhanced muscle-specific transfer by AAVs. **a**, Diagram for systemic treatment of DCC mice. Mice (aged 6-8 weeks) were injected with PBS, AAV9-ARC-S, or AAV9-ARC-T. Blood was collected at 6 and 12 weeks after injection, and CD4^+^ and CD8^+^ T cells were purified from PBMCs to assess immunogenicity (n=6). **b,c,** Immunogenicity analysis for T cells response. Left, flow cytometry analysis of CD4^+^ (b) and CD8^+^ (c) T cells; Right, cytokine levels at 6 and 12 weeks after injection. Relative expression levels were normalized to the PBS-treated mice. Data are mean ± s.e.m.(n=6). **d,** The expression levels of ARCs in various tissues after intravenous injection. Data were normalized to the ARC expressions in the heart. Center line, median; box limits, upper and lower quartiles; whiskers, min-max (n=6). **e,** Relative expression levels of CRE and non-CRE RNAs in different tissues of ARCs-treated mice. Data are mean ± s.e.m. (n=6). **f,** Representative fluorescence imaging of tissue tropism of different AAV serotypes. Mice were injected with either AAV9-tdTomato or iAAV9.6-tdTomato (4×10^13^ vg/kg) *via* tail vein at 4 weeks old. The liver sample was exposed for 200 ms, while all other tissues were exposed for 600 ms. An extra intensity gain (300%) was used for muscle tissues. Scale bar, 500 μm. Statistical significance was assessed by an unpaired two-tailed Student’s *t*-test (\*\**P*< 0.01; \*\*\**P*< 0.001; \*\*\*\**P*< 0.0001).

We further examined the expression of ARCs in various tissues after the systematic delivery by AAV2/9, and found that the ARCs were predominantly expressed in the liver and heart, with low expression levels in muscle tissues (Fig. 4d). As a result, the toxic RNA repeats were insufficiently reduced in the muscle tissues compared to the liver (Fig. 4e). These results also reflected the current limitations of delivery technologies ^53,54^. To improve muscle delivery efficiency, we applied a new generation of AAV vector, iAAV9.6 (patent application CN202510458749.9), which preferably infect muscle tissues. In a head-to-head comparison using tdTomato as a reporter gene, iAAV9.6 showed greater delivery capacity for cardiac and major skeletal muscles compared to regular AAV9 (Fig. 4f). Importantly, the myotropic iAAV9.6 exhibited lower liver tropism than AAV9, suggesting reduced hepatotoxicity, enhanced safety, and strong potential as a gene therapy vector for muscle disorders (Fig. 4f).

### Systemic treatment with AAV-ARCs in HSA_LR_ mouse model

To further tested the *in vivo* efficacy of ARCs in restoring muscle defect of DM1, we used iAAV9.6 to systematically deliver the ARCs into HSA_LR_ mice that exhibited many of the characteristics of myotonic dystrophy ^55,56^ (Fig. 5a). To enhance ARC expressions, we optimized ARC codons with different computational tools (Supplementary Table 2) and tested the expression of the ARCs (Extended Data Fig. 8). Additionally, we used human muscle creatine kinase promoter to drive expression of ARC-S and ARC-T in muscles ^57^. We confirmed that the muscle-specific iAAV9.6 can effectively deliver ARC-S to various muscle tissues and heart with much higher expression than other non-muscular tissues (Fig. 5b). Compared to PBS-injected mice, we observed a 40%-60% reduction in pathogenic RNA levels induced by ARC-S across multiple muscle types (Fig. 5c). In addition, RNA-seq analysis showed the ARC-S treatment had minimal impact on global gene expression (22 genes downregulated and 30 genes upregulated), including the CUG-containing genes (Extended Data Fig. 9a and 9b). Moreover, the affected genes were closely associated with cell junction organization or muscle system (Extended Data Fig. 9c). Meanwhile, the mis-regulated expression of muscle-related genes were also rescued by ARC-S treatment (Extended Data Fig. 9d).

**Fig. 5.**
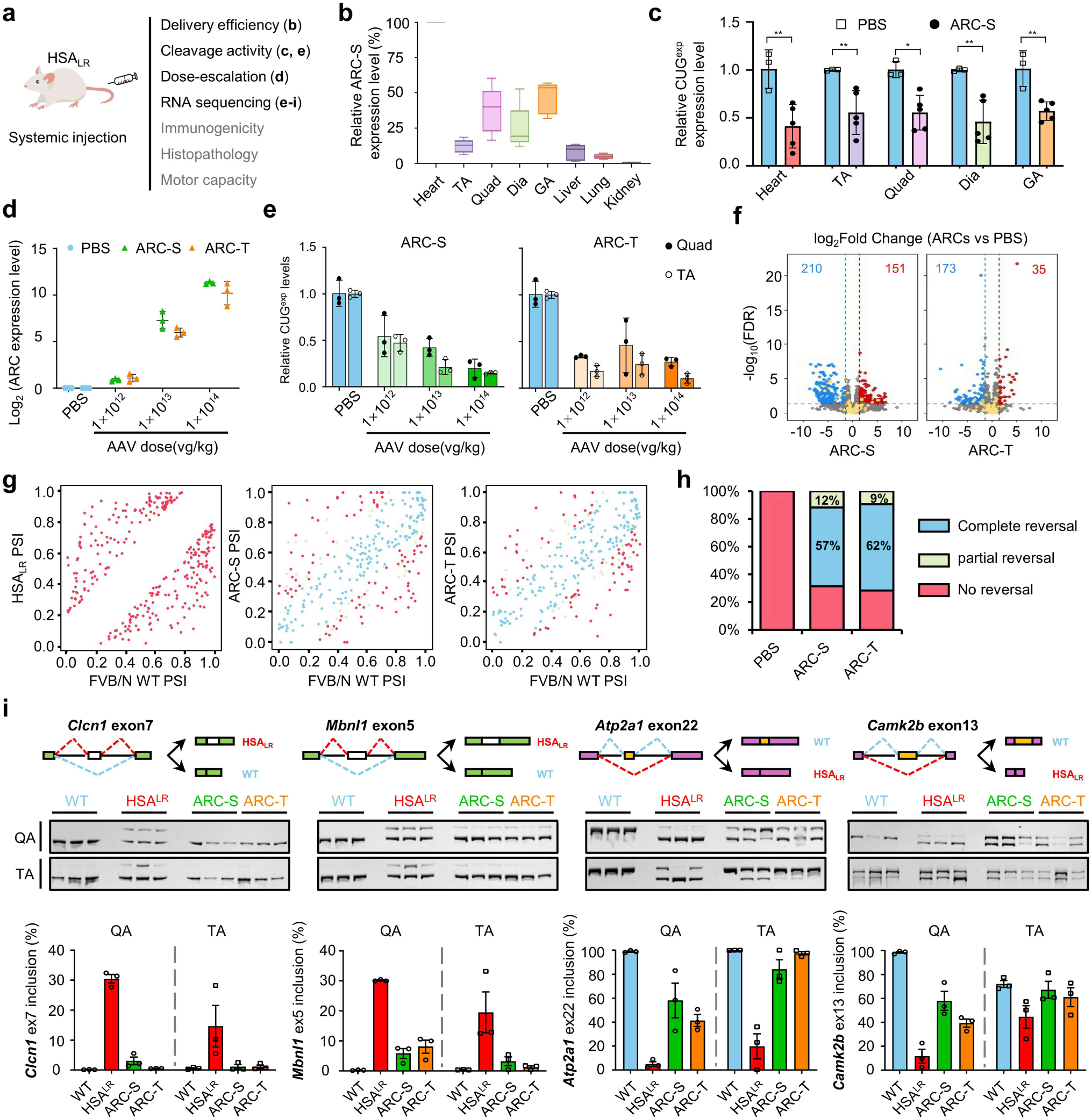
Systemic treatment with iAAV9.6-ARCs corrects pathogenic defects in HSA_LR_ mice. **a**, Schematic for experimental design. The HSA_LR_ mice (aged 6-8 weeks) were injected with PBS or iAAV9.6-ARCs driven by a muscle-specific promoter. Samples were collected at 8 weeks post-injection for subsequent analyses. **b,** The expression levels of ARC-S in various tissues after intravenous injection. Data were normalized against ARC-S expression in the heart. TA: Tibialis anterior; Quad: Quadriceps; Dia: Diaphragm; GA: Gastrocnemius. Center line, median; box limits, upper and lower quartiles; whiskers, min-max (n=5). **c,** Levels of CRE transcripts determined by RT-qPCR in multiple muscles tissues of HSA_LR_ mice. PBS or iAAV9.6-ARC-S (1×10^12^ vg per mouse) was delivered into HSA_LR_ mice (aged 8 weeks) by intravenous injection. Data are mean ± s.d. (n = 3 for PBS control; n=5 for ARC-S treated group). **d,** The expression levels of ARCs with varying AAV doses. Data are mean ± s.d. (n = 3). **e,** Levels of CRE transcripts in Quad and TA of AAV-treated mice. Data are mean ± s.d. (n = 3). **f,** Global analyses of gene expression in treated tibialis anterior of HSA_LR_ mice. Yellow dots indicate genes with ≥3 CUG repeats, while red and blue dots represent up- and down-regulated genes (adjusted *P* < 0.05, |log_2_FC| > 1). **g,** Scatter plot of transcriptomic analyses shows overall correction of splicing defects (HSA_LR_/treated mice *vs.* wild type mice). Red dots indicate the mis-splicing events; blue dots indicate the complete reversal events; green dots indicate the partial reversal events (FDR < 0.05, |Δ PSI|> 0.2, n= 3). **h,** Summary of mis-spliced AS events reversed by ARC-S and ARC-T. **i,** Representative AS defects corrected by ARCs. RT-PCR analysis revealed mis-splicing events in *Clcn1* exon 7, *Mbnl1* exon 5, *Atp2a1* exon 22, and *Camk2b* exon 5 in Quad) and TA muscles of treated HSA_LR_ mice (AAV, 1×10^14^ vg/kg), compared to PBS-injected and wild-type controls. Data are mean ± s.e.m. (n=3). Statistical significances were performed by the unpaired two-tailed Student’s *t*-test (\**P*< 0.05; \*\**P*< 0.01).

Encouraged by these results, we conducted a comprehensive dose-escalation study of ARCs in HSA_LR_ mice, followed by multifaceted analyses of phenotypes (Fig. 5a). We administered iAAV9.6-ARCs with a dosage gradient (1×10^12^, 1×10^13^, and 1×10^14^ vg/kg) in HSA_LR_ mice and observed sustained tolerability of the treated mice across the given doses, with no mortality cases. As expected, the expression level of both ARCs increased with dose escalation (Fig. 5d). Eight weeks post-injection, the expression levels of aberrant CREs decreased by >50% in the tibialis anterior (TA) and quadriceps (QA) of the treated mice, with up to ∼80% reduction in the high-dose groups (Fig. 5e).

The subsequent transcriptomic analysis of the high dose groups revealed that approximately 1-2% of the genes were affected in the ARC-treated HSA_LR_ mice (361 genes for ARC-S; 208 genes for ARC-T) (Fig. 5f). Global dysregulation in RNA alternative splicing is a typical feature of DM1, which can serve as a key criterion in evaluating potential therapeutics ^56^. We found the mis-regulated exon-skipping events in HSA_LR_ mice could be efficiently restored by ARC-S and ARC-T (Fig. 5g). Overall, more than 70% of mis-splicing events were reversed in TA (Fig. 5h; ARC-S: 57% complete reversal and 12% partial reversal; ARC-T: 62% complete reversal and 10% partial reversal). Gene ontology analysis revealed that ∼300 altered AS events were significantly enriched in muscle functions and regulation of RNA metabolism, indicating the molecular pathology of DM1 (Extended Data Fig. 10). Sequestration of MBNL1 leads to pathological splicing alterations ^58^, such as the inclusion of exon 7a in the chloride voltage-gated channel 1 (*Clcn1*), a hallmark of DM1 that contributes to myotonia ^59^. The quantitative measurement for the representative mis-spliced exons in DM1 (exon 7a in *Clcn1*, exon 22 in *Atp2a1*, exon 5 in *Mbnl1* and exon 13 in *Camk2b*) confirmed that splicing defects had been efficiently corrected in QA and TA treated by ARCs (Fig. 5i).

### Systemic treatment with AAV-ARCs alleviated the pathologic phenotype of DM1

We further evaluated the therapeutic performance of ARCs at both muscular pathology and motor capacity in HSA_LR_ mice. As expected, using histological staining and RNA-FISH, we monitored increased number of peripheral muscle nuclei and decreased RNA foci in the TA of HSA_LR_ mice (Fig.6a, 6b and Extended Data Fig. 11), a pathologic feature reported previously ^55^. However, in the ARCs-treated groups, we observed a progressive change of peripheral muscle nuclei and RNA foci in an AAV dose-dependent manner (Fig.6a and 6b), suggesting a mitigation of morphological phenotype.

To assess the motor coordination and endurance of mice, we conducted the rotarod test for all experimental groups (Fig.6c), a behavioral test that is commonly used to measure muscle stamina. Compared to PBS-treated HSA_LR_ mice, we found that the ARCs-treated mice exhibited prolonged movement duration and extended travel distance (Fig.6c). In the high-dose groups, the endurance and mobility of both ARC-S and ARC-T treated groups were close to the wild-type mice. In addition, the myotonia of HSA_LR_ mice was evaluated by the latency between hindlimb flexion and relaxation (Fig. 6d). The hindlimb pull-test showed that PBS-treated mice displayed classic signs of myotonia, including prolonged muscle stiffness and delayed relaxation after stretching. In contrast, ARCs-treated mice rapidly regained muscle relaxation within a significantly shorter latency than the PBS-treated group, with the high-dose ARC-S group close to the wild type (Fig. 6d, Supplementary Video 1 and 2).

**Fig. 6.**
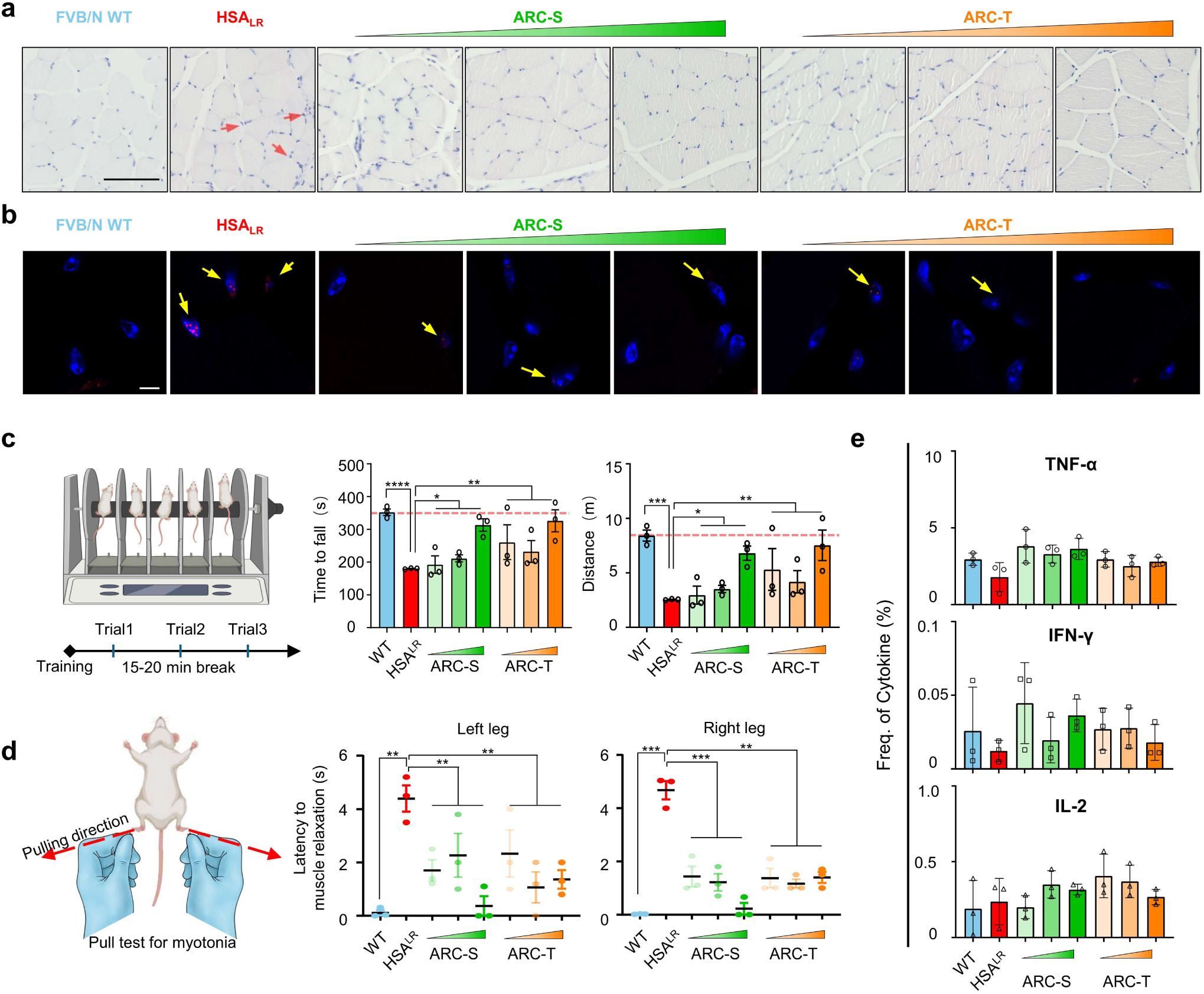
Systemic treatment with iAAV9.6-ARCs rescues DM1-related phenotypes. **a**, Representative H&E staining of tibialis anterior from ARCs or PBS treated HSA_LR_ mice. The arrowheads highlight the presence of peripheral muscle nuclei in HSA_LR_ mice (red). Scale bar, 200 µm. **b,** RNA-FISH analysis of CUG repeats (red) and cellular nucleus (blue) of latitudinal sections of tibialis anterior treated with ARCs (representative image, n = 3). Blue, DAPI. The arrowheads highlight the presence of CUG RNA foci (yellow). Scale bars, 5 µm. **c,** Schematic illustration of rotarod test. ARCs or PBS treated HSA_LR_ mice (aged 14 weeks) and WT mice were pre-trained and then subjected to three rotarod trials with a rest interval between each trial. The quantification results of falling time (left) and total distance (right) were plotted. Data are mean ± s.e.m. (n=3). **d,** The schematic and quantification of mice pull-test. Relaxation time of latency to hind-limb was recorded for the left and right legs separately. Data are mean ± s.e.m. (n=3). **e,** Immunogenicity analysis for T cells response. Cytokine measurement for iAAV9.6-ARCs at 8 weeks post-injection. Frequency of TNF-α, IFN-γ and IL-2 detected by flow cytometry, with dosage indicated (n= 3). Statistical significance was assessed by an unpaired two-tailed Student’s *t*-test with Welch’s correction (\**P*< 0.05; \*\**P*< 0.01; \*\*\**P*< 0.001).

The immune response induced by exogenous proteins is the major concern of gene therapy, and thus we also measured the possible T cells responses triggered by injected iAAV9.6-ARCs. Consistent with our previous findings, the cytokine levels of TNF-α, IFN-γ and IL-2 showed minimal changes (Fig. 6e), suggesting that ARCs represent a safe therapeutic approach with low immunogenicity for treating CUG repeat expansion diseases. Collectively, our comprehensive studies using a gold-standard DM1 mice model demonstrated that the systematic administration of ARCs can restore the aberrant molecular pathology and major muscle functions without causing detectable immune response, highlighting the therapeutic potential of ARCs as an effective and safe treatment for DM1.

## DISCUSSION

DM1 is a multi-system disorder, with age of onset and severity determined by the CRE lengths. A variety of therapeutic strategies based on small RNA and CRISPR/Cas have been developed to target the abnormal CREs, aiming to alleviate or reverse the molecular phenotype in DM1. However, there is currently no approved therapy with proven safety, efficacy, and durability, and the development of new DM1 therapy remains a major challenge. In this study, we engineered a series of ARCs and screened their activities using a newly developed dual-fluorescence reporter. The resulting ARC-S and ARC-T were further optimized to specifically and efficiently degrade CREs in a length dependent fashion. The efficacy, off-target effects and toxic side-effects were further tested in DM1 animal models, and the results showed that both ARCs can effectively degrade toxic RNA repeats and rescue DM phenotypes without causing detectable immune response, indicating their potential as a promising avenue for gene therapy of diseases caused by toxic RNA repeats.

Safety concerns in gene therapy primarily revolve around the immunogenicity of the drug and the off-target effects of the treatment. To minimize the immunogenicity of ARCs, we constructed and screened ARC components with human-originated proteins (rather than bacterial proteins), thereby reducing the potential need for a life-long immunosuppressive treatment ^34^. Consistent with this idea, the systemic administration of ARCs in DCC and HSA_LR_ mouse models revealed no significant activation of CD4^+^ or CD8^+^ T cells, indicating minimal immune response (Fig 4b,4c and 6e). Furthermore, histological analysis showed no pathological changes or lesions in muscle tissues (Fig 6a). Our strategy to reduce immunogenicity has been employed in another study using engineered MBNL1Δ proteins ^21^. The low immunogenicity of ARCs may also enable repeated administration with different AAV variants. In contrast, non-human proteins are more likely to provoke immune responses, thereby limiting their therapeutic efficacy and long-term effectiveness. For example, the state-of-the-art therapeutic methods for DM1 using bacterial Cas proteins (PIN-RCas9 system and Cas13d) require additional immunosuppressive treatment ^34^, complicating repeated administration and limiting clinical applicability (Supplementary Table 1). Additionally, the ARC system functions at the RNA level by selectively degrading pathogenic CREs without altering the genomic DNAs. Through transcriptome-wide studies in cells and animal models, we confirmed that ARCs can effectively reverse CREs-induced alterations of transcriptome with lower off-target rate. The newly developed iAAV9.6 facilitates targeted delivery to muscle tissue while preventing liver accumulation (Fig 4f and 5b), thereby enhancing therapeutic safety. Collectively, our results showed that the ARC system delivered *via* the tissue-specific AAV capsid may provide a promising therapeutic platform with superior safety.

The therapeutic efficacy of ARCs is supported by evidence on two levels: at the molecular level, ARCs reduce intranuclear RNA-protein granules and release sequestered RNA-binding proteins; at the phenotypic level, ARCs restore muscle function and improve motor ability. The imaging and RNA-seq analyses in animal models suggested that ARCs preferably reduce the level of abnormal CREs while showing limited effect on the normal CUG-containing transcripts, which is a key advantage over existing methods. Although both short and long CUG repeats can theoretically be recognized by ARCs, the longer repeats have more binding sites that probably increase the probability for ARCs recruitment and CREs degradation. In addition, the CREs are primarily “trapped” in the nucleus, while the normal transcripts containing short (CUG)_n_ repeats are largely transported into cytoplasm. In our design, we included multiple NLS to ensure the nuclear localization of ARCs, which further contributed to their preference in degrading longer CREs in the nucleus.

Despite recent successes with AAV-based systems, efficient delivery remains a major challenge in gene therapy. The compact size of the ARC system (∼2 kb) enables its delivery with a single AAV vector, whereas the CRISPR/Cas-based approaches often exceed the packing capacity of AAV (PIN-RCas9, ∼4.7kb;Cas13d, ∼2.9kb;dSaCas9, ∼3.1kb) and thus may require dual-vector systems (Supplementary Table 1). In addition, consistent with previous reports ^60–62^, the wild-type AAV9 showed limited delivery efficiency for skeletal muscle (e.g., tibialis anterior, quadriceps and heart) (Fig.4d). To solve this problem, we employed muscle-tropic capsids iAAV9.6 (patent application: CN202510458749.9) that showed strong and widespread expression in skeletal muscle with minimal off-target activity. Dose-escalation study suggested that the iAAV9.6 can mediate dose-dependent expression of ARCs in muscles (Fig.5d), achieving significant degradation of pathogenic CREs even in low doses (Fig.5e). Importantly, intravenous AAV doses used in FDA-approved gene therapy trials typically range from 1-3×10^14^ vg/kg ^60,62^. In our study, all the effective ARC doses (1×10^12^-1×10^14^ vg/kg) were within this established safety range, indicating that iAAV9.6-delivered ARCs can achieve precise muscle targeting and strong therapeutic effects at clinically acceptable doses.

The ongoing clinical trials of DM1 focus on enhancing small RNA or ASO delivery efficiency by coupling with polypeptides (e.g., NCT06667453, PepGen Inc) or antibodies (e.g., NCT06411288, Avidity Biosciences; NCT05481879, Dyne Therapeutics). These approaches often require frequent administration. However, because muscle comprises up to 40% of total body weight, systematic therapies demand extremely high dosage for muscular diseases, increasing the risk of dose-related toxicity. The iAAV9.6 enabled efficient and durable treatment in DM1 mouse model. Practically, therapeutic effects could be sustained through redosing with another AAV serotypes, supporting long-term clinical benefit.

Our comprehensive study provides the proof-of-concept for AAV-mediated DM1 therapy by ARCs, demonstrating effective phenotype reversal in HSA_LR_ mouse models. However, as human DM1 patients may exhibit larger pathological expansions (up to 10000 repeats), evaluating the therapy in models with various repeats would provide better assessment of its efficacy and limitations. Extensive preclinical tests, including dose escalation experiments, biodistribution, and immunogenicity evaluations, were performed in this study to establish the clinical relevance of the ARC system. The results suggest that the ARC system meets rigorous preclinical standards and shows great promise as a clinically viable treatment for DM1. Future studies in non-human primates will be conducive to clarify ARC’s safety and efficacy across different doses.

---

**Extended Data Fig. 1.**
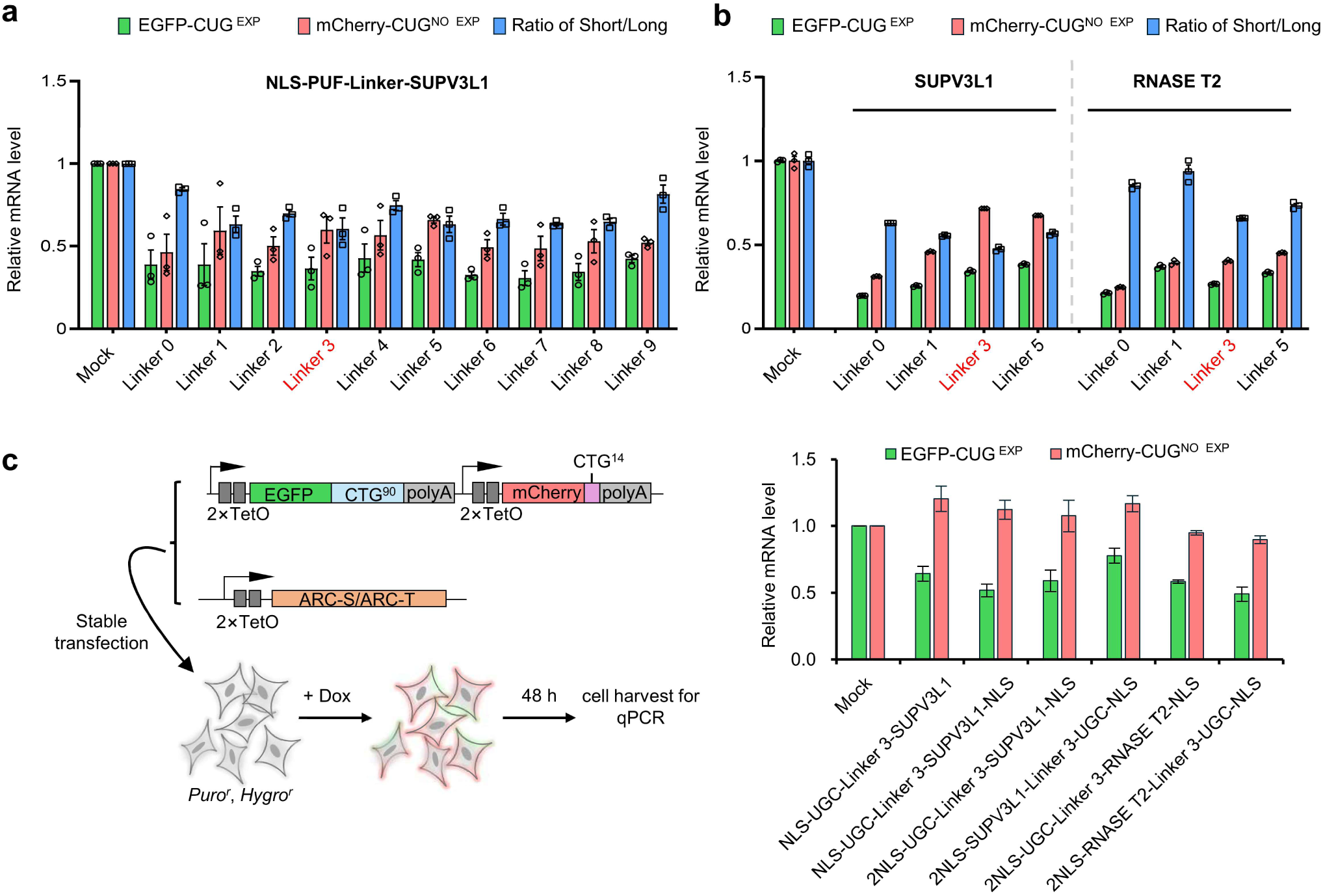
Optimization and validation of the ARCs for CRE degradation. **a**, Linker optimization for ACRs. Linkers refers to the 7-11 amino acid peptide selected from the Linker database (http://www.ibi.vu.nl/programs/linkerdbwww/). Relative EGFP and mCherry mRNA levels in NLS-UGC-SUPV3L1 (with various Linkers) transfected cells were normalized to mock. The original linker was noted as Linker 0. Data are mean ± s.d.(n=3). **b,** Linker 1, 3, and 5 were selected and tested by fusing with RNase domains of SUPV3L1 and RNASE T2 to determine the optimal linker. The original linker 0 was included as a control. Data are mean ± s.d.(n=3). **c,** Validation of the optimal ARC constructions in the stable cell line. The dual-fluorescence reporter and ARCs constructs were co-transfected and then the cells were selected using Hygromycin (*Hygro*^r^) and Puromycin (*Puro*^r^) resistance, respectively. Quantification of the CRE and non-CRE RNA abundance in *Hygro*^r^ and *Puro*^r^ positive cells (bottom). TetO, tetracycline operator. Dox, Doxycycline. Data are mean ± s.d.(n=3).

**Extended Data Fig. 2.**
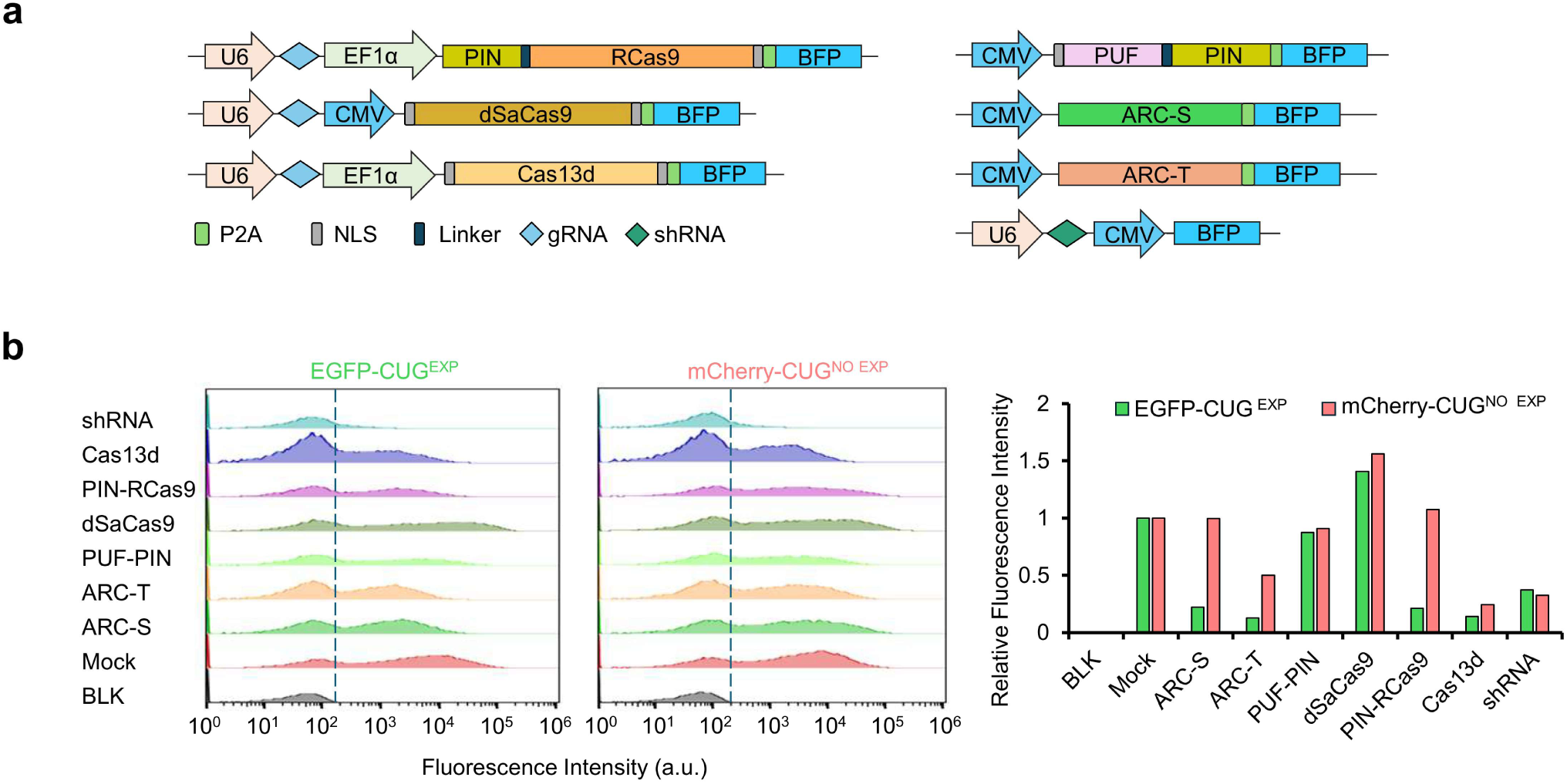
Characterization of the efficiency and specificity of current CRE-targeting tools. **a**, Schematic diagraphs for various CRE-targeting tools. NLS, nuclear localization signal. P2A, 2A peptide from porcine teschovirus-1. **b,** Relative fluorescence intensity detected by flow cytometry. Histograms showed the fluorescence signals in HEK 293T cells co-transfected with the dual-fluorescence reporter and various CRE-targeting tools (left). The fluorescence intensities were normalized to the mock group. > 20000 Cells were counted for each group.

**Extended Data Fig. 3.**
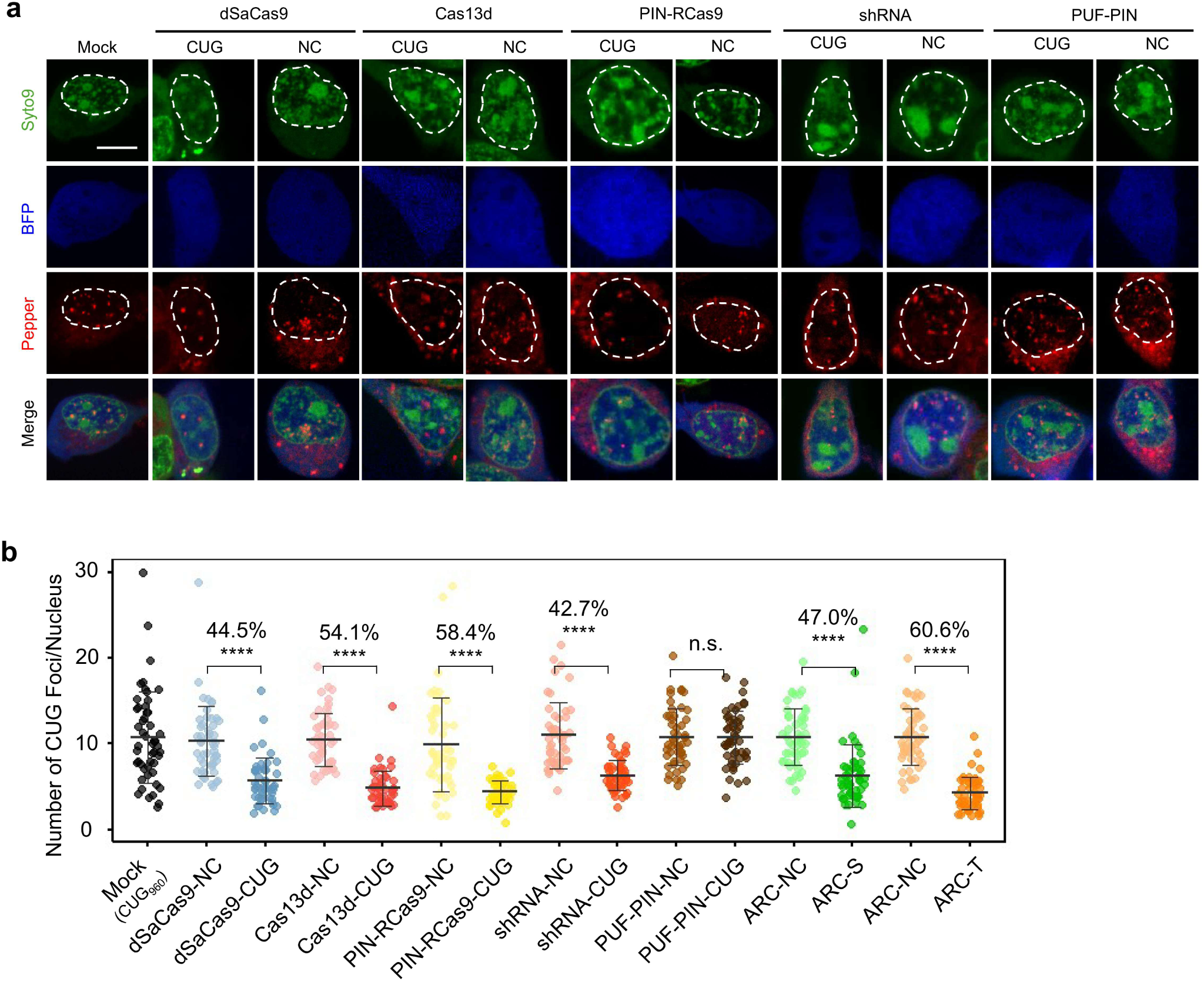
Comparison of current CRE-targeting tools in eliminating RNA foci. **a**, Live-cell image of nuclear RNA foci. HEK 293T cells were co-transfected with the BFP tagged CRE-targeting tools and *DMPK* reporter containing 960 CTG repeat tracts. The RNA foci were indicated by the Pepper (red). Syto9 staining indicates the nuclear region (green), and blue indicated localization of BFP-tagged ARCs (blue). Scale bar, 5 μm. **b,** Statistical analysis of the RNA foci number. Comparison of the RNA foci degradation efficiency. Data represents the mean ± s.d. (n = 50 cells). The numbers above the lines represented degradation efficiency. Statistical significances were performed by the two-tailed Student’s *t-*test (\*\*\*\**P*< 0.0001; n.s., not significant).

**Extended Data Fig. 4.**
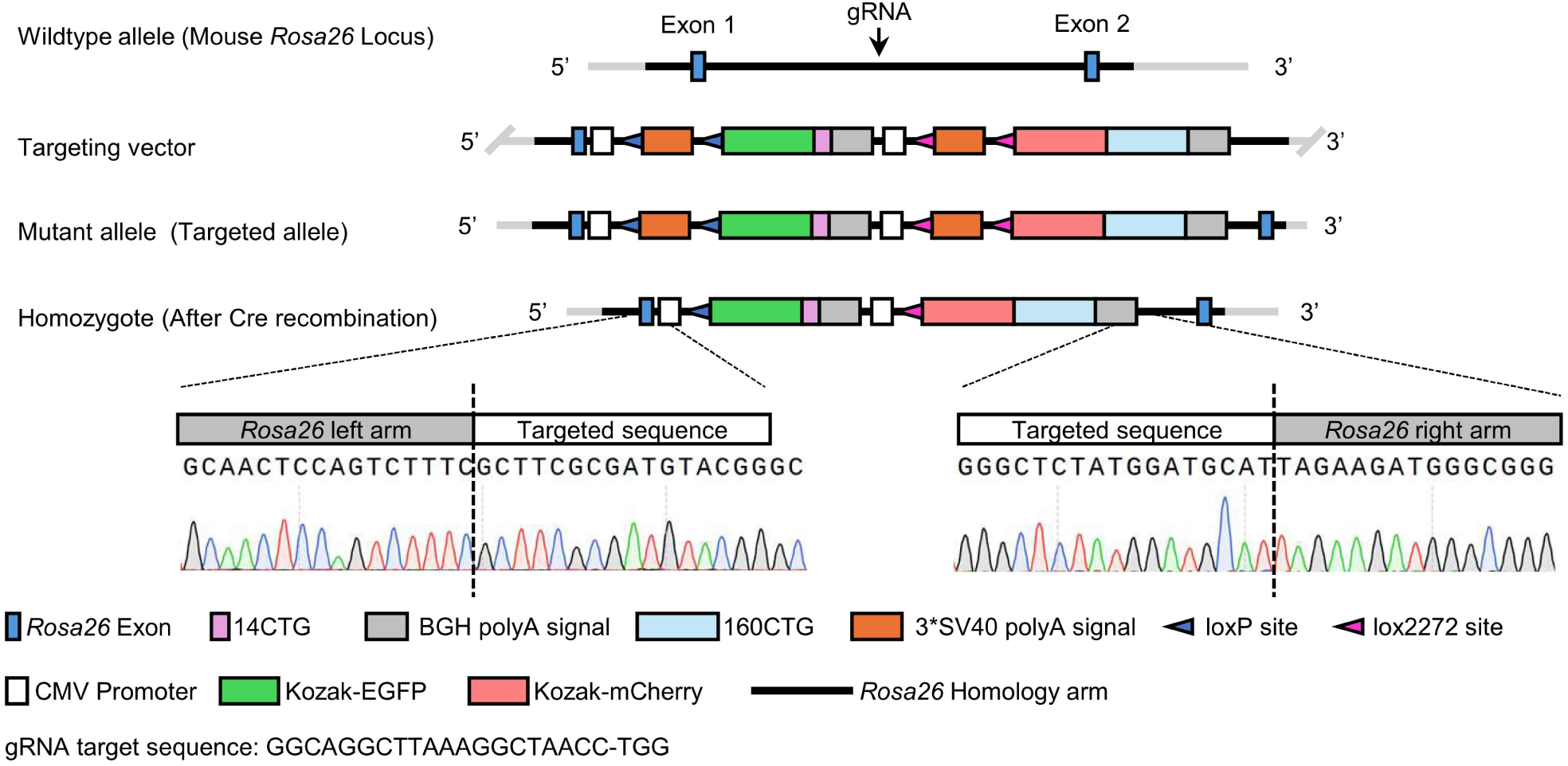
The construction strategy of the DCC transgenic mouse model. Genetic design of the <u>D</u>ual <u>C</u>olor <u>C</u>UG^exp^ (DCC) transgenic mouse model. Dual-fluorescence reporters were integrated into the endogenous *Rosa26* locus of C57BL/6 mice by CRISPR-Cas9. The homozygosity was confirmed by Sanger sequencing of flanking regions after Cre-mediated recombination.

**Extended Data Fig. 5.**
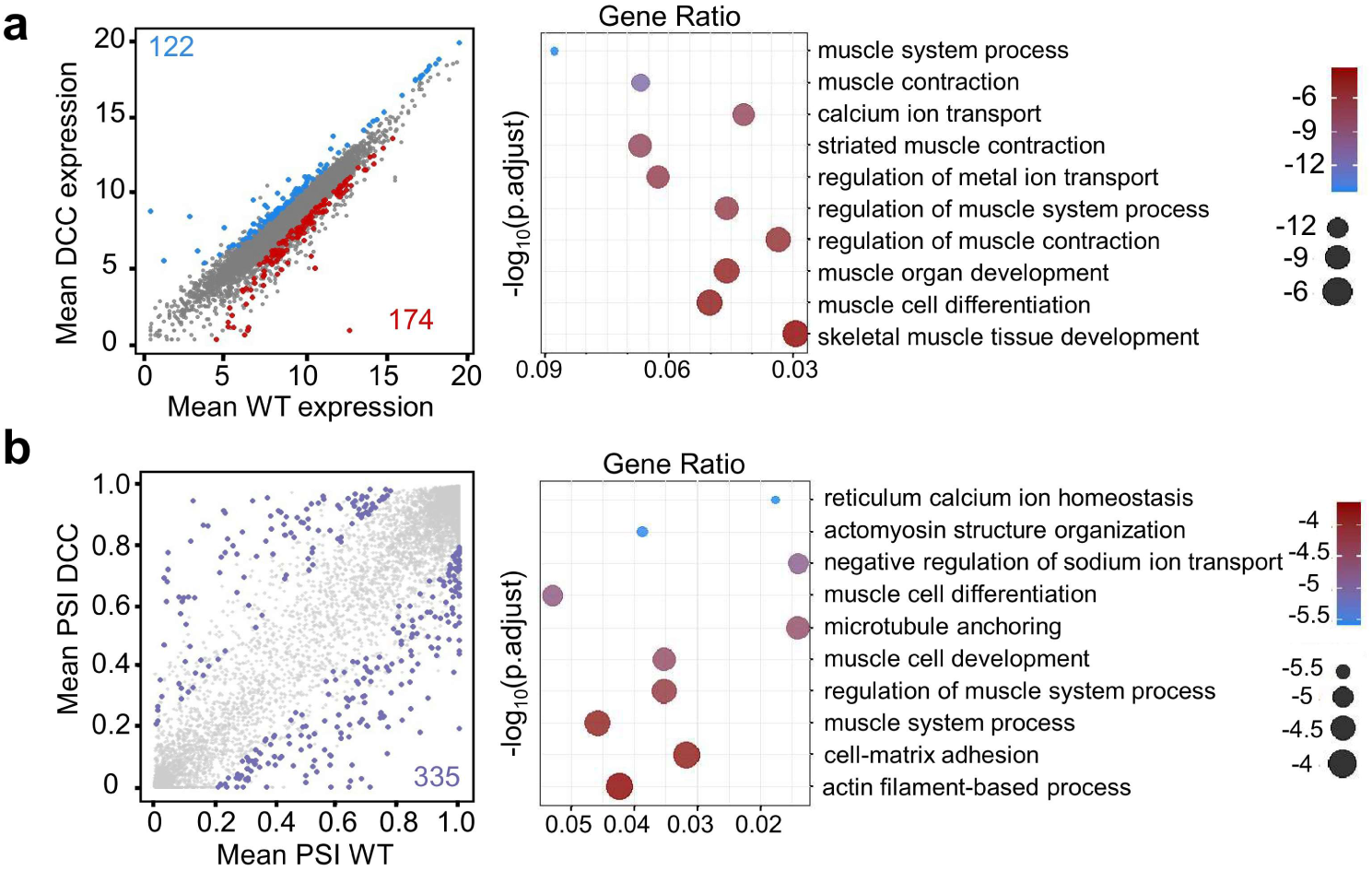
Transcriptome-wide analysis of the DCC transgenic mouse model. **a-b**, Transcriptome-wide analyses of the global gene expression (**a**) and alternative splicing (**b**) between DCC and WT mice. The red dots represent up-regulated genes, and the blue dots represent down-regulated genes (adjusted *P*< 0.05, |log_2_FC| >1). The purple dots represent altered AS events (FDR < 0.05, |Δ PSI|> 0.2). The GO analysis of differentially expressed genes and splicing events were performed, with the top 10 related GO terms plotted on the right. *P* values were calculated using modified Fisher’s exact test (n= 3).

**Extended Data Fig. 6.**
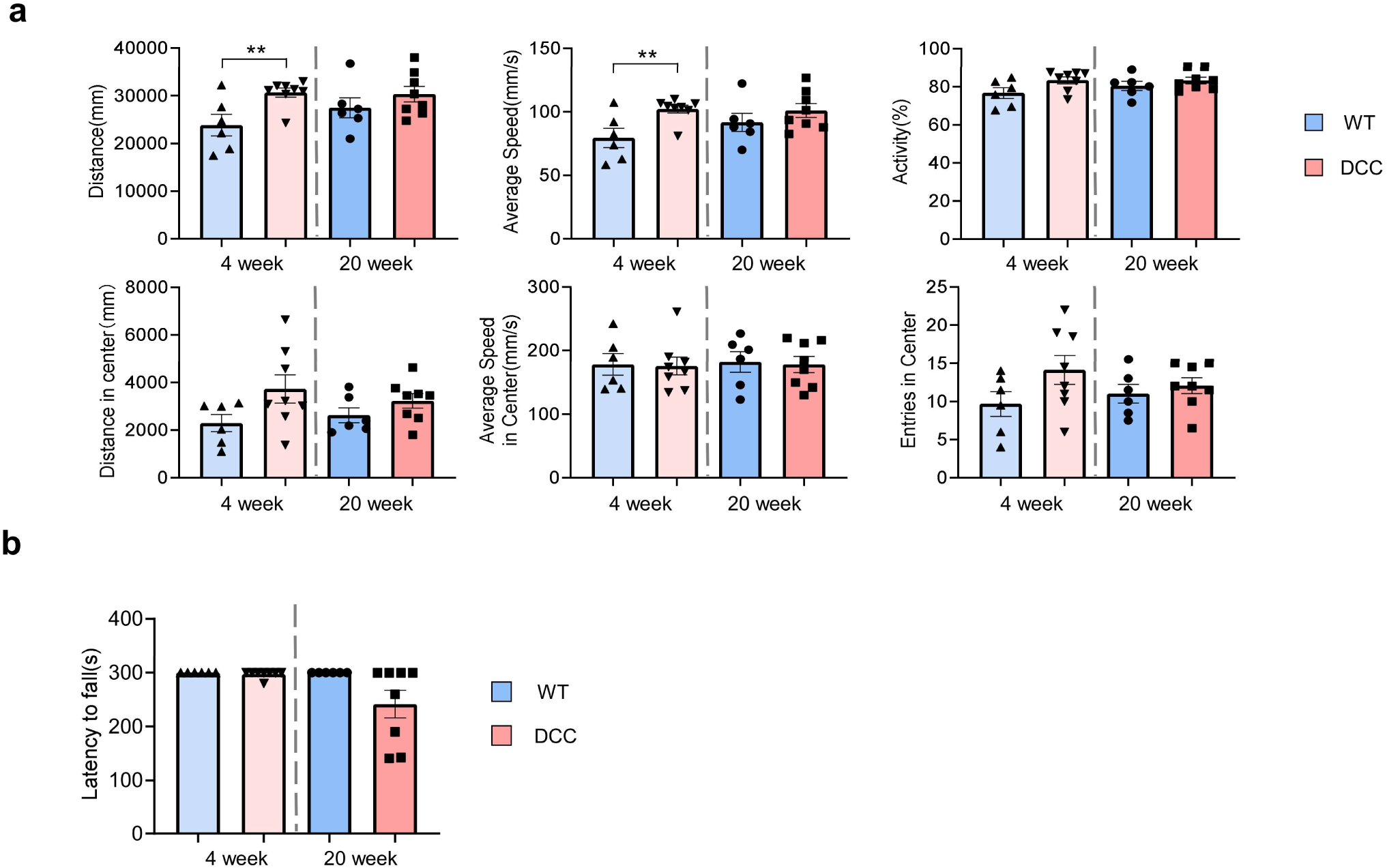
Behavioral test for DCC and wild-type mice. **a**, Open field test of DCC and wild-type mice. Distance, average speed, activity, and corresponding center area indices were measured at 4 and 20 weeks of age (n=6-8). **b,** Rotarod test of DCC and wild-type mice. Rotarod falling time was recorded at 4 and 20 weeks of age (n=6). Statistical significances were performed by an unpaired two-tailed Student’s *t-*test (\*\**P*< 0.01).

**Extended Data Fig. 7.**
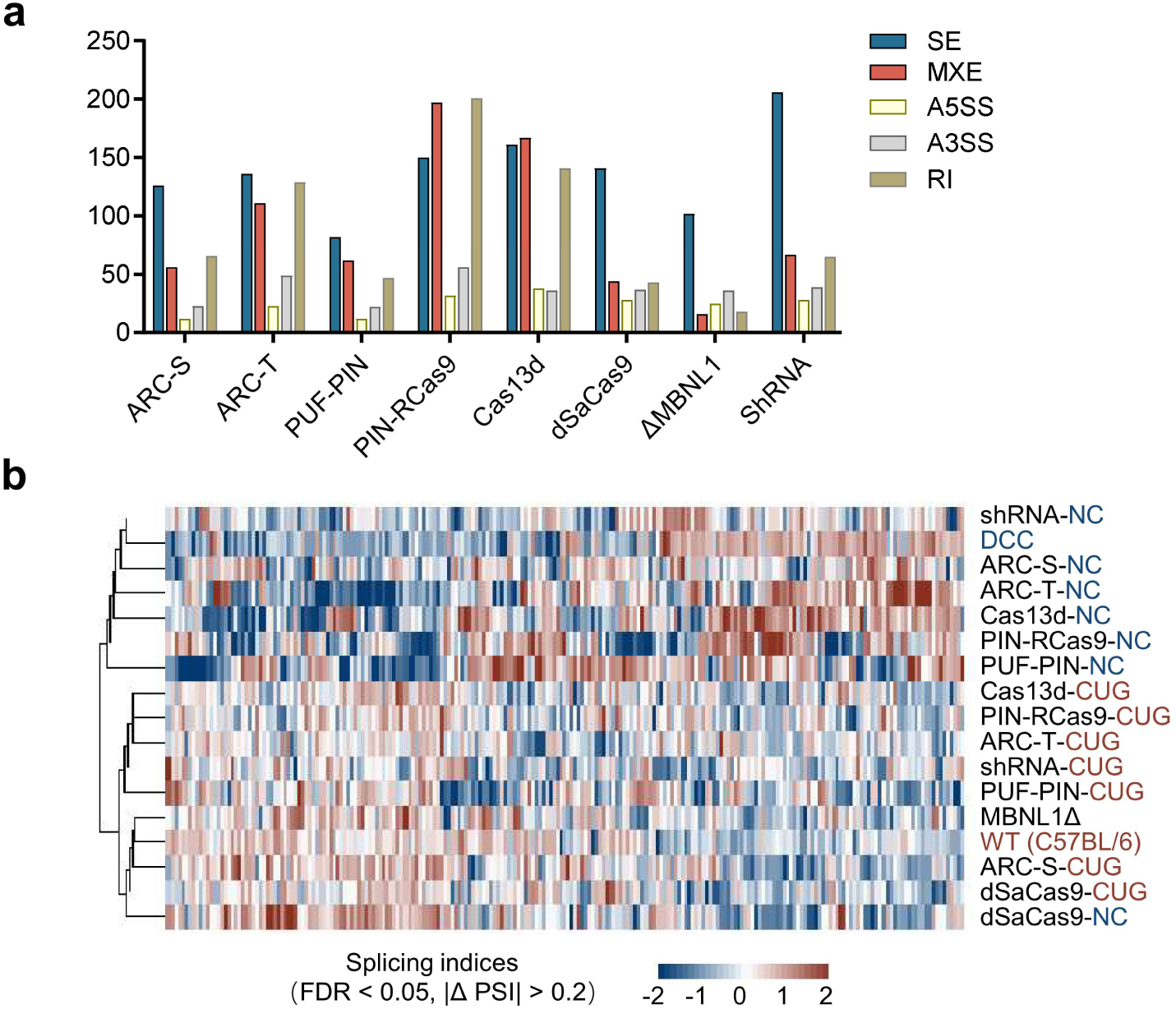
Splicing alternation in AAV-treated quadriceps of DCC mice. **a**, Numbers of alternative splicing events in different experimental groups.SE, skipped exon; MXE mutually exclusive; A3’SS/A5’SS alternative 3’/5’ splice sites; RI, retained intron. **b,** Hierarchical clustered heatmap of exon inclusion indices in AAV-treated quadriceps muscle (Targeting *vs.* Non-targeting) (n=3 for each group). The CB57L/6 wild type mice and PBS-treated DCC mice were highlighted.

**Extended Data Fig. 8.**
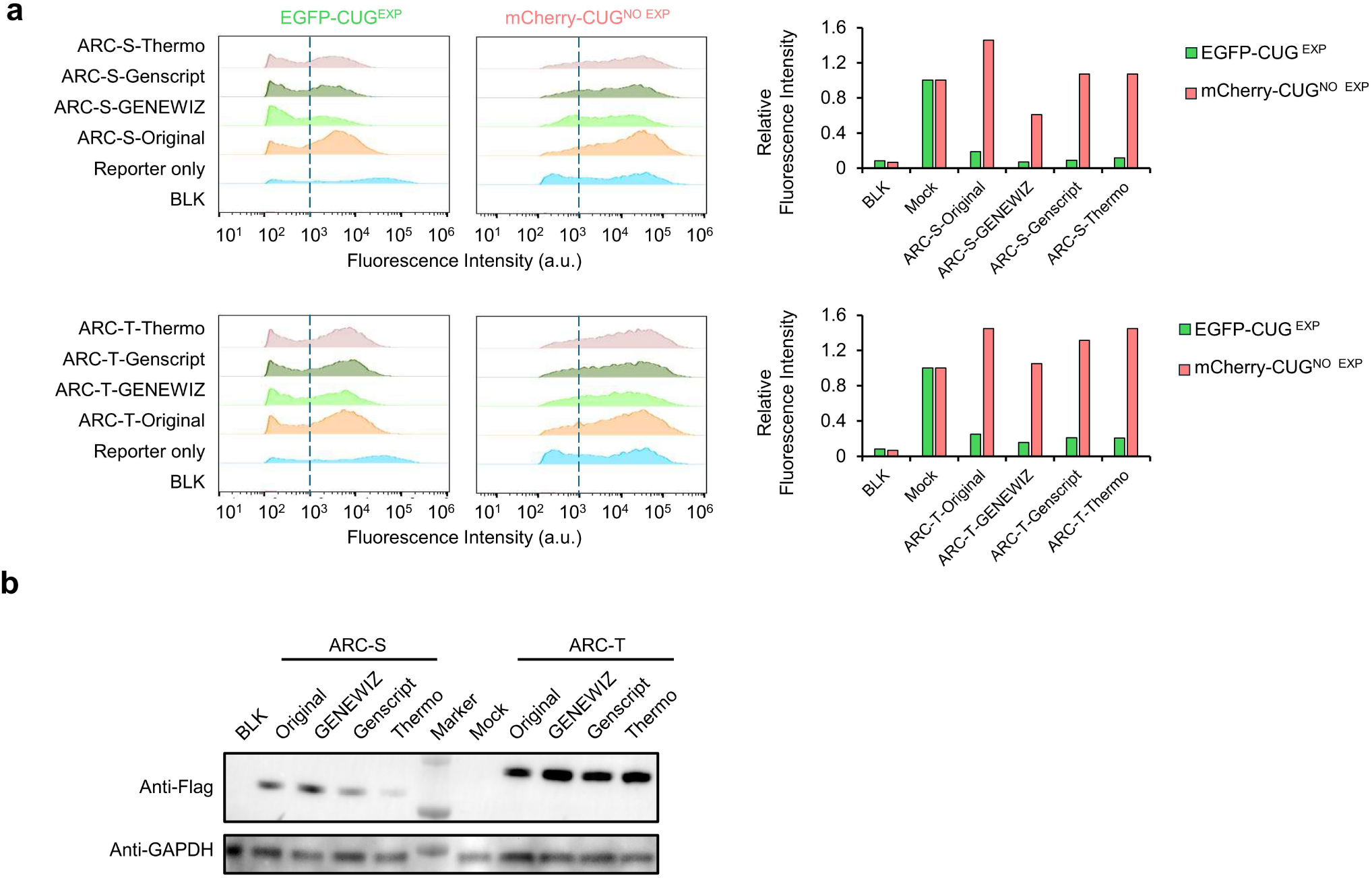
Codon optimization of ARCs to enhance their performance. **a**, Dual-fluorescence reporter assay to evaluate the enhancement of the codon optimized ARCs. ARCs were optimized by three online tools (GENEWIZ, Inc., GenScript Biotech and Thermo Fisher Scientific Inc.). Histograms show fluorescence changes in HEK 293T cells co-transfected with the reporters and optimized ARCs (left); relative fluorescence intensity for each ARCs normalized to Mock group (right). **b,** Western blot analysis of various optimized ARCs, with GAPDH used as a loading control.

**Extended Data Fig. 9.**
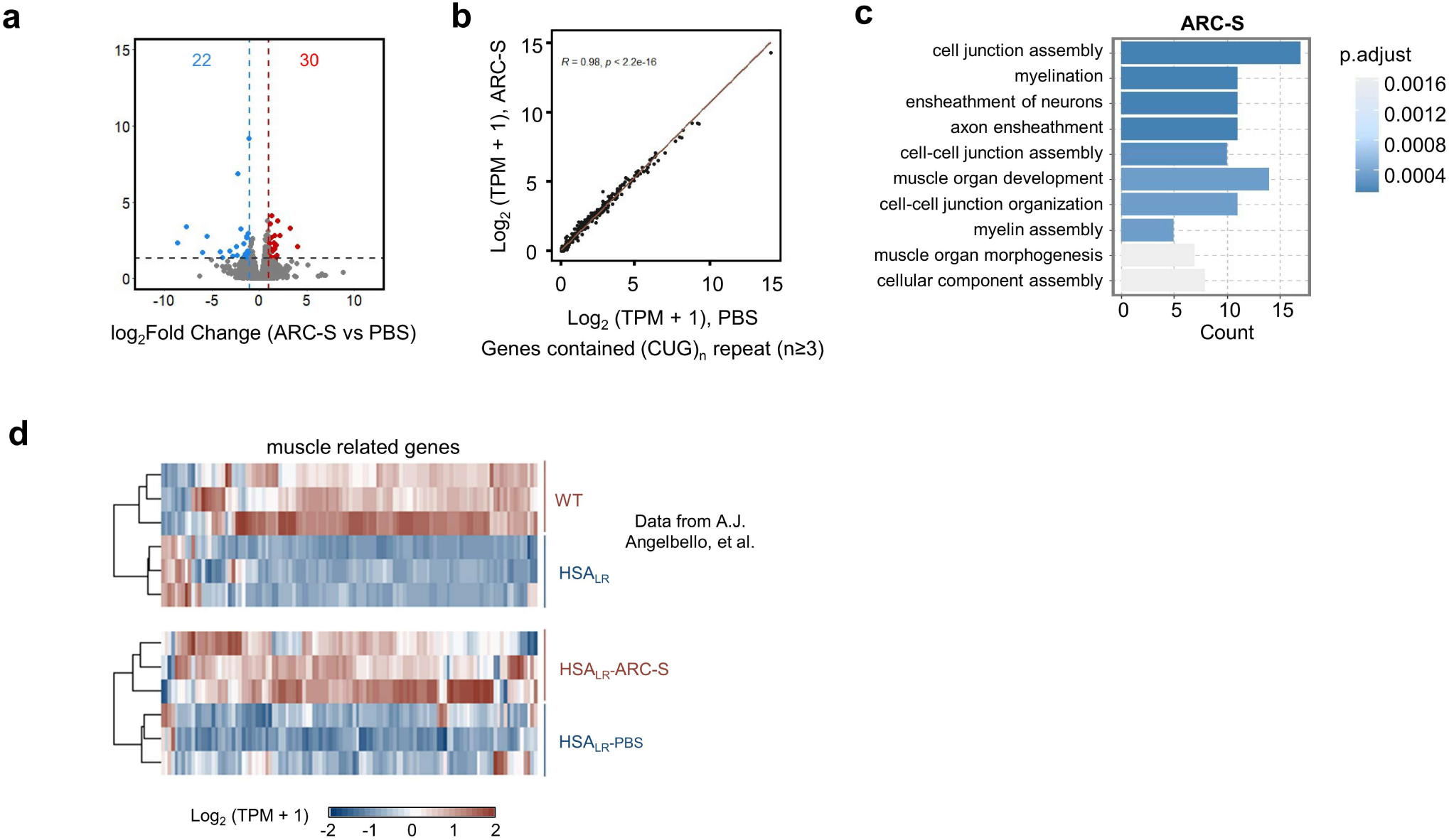
Transcriptome-wide analysis of gene expression in the tibialis anterior of HSA_LR_ mice. **a**, Volcano plot illustrates global gene expression changes in tibialis anterior (TA) of ARC-S treated-HSA_LR_ mice after 8 weeks post-injection. The number of altered genes were indicated above. Red and blue dots represent up- and down-regulated genes in ARC treated mice (adjusted *P*< 0.05, |log_2_FC| >1). **b,** Scatterplot shows expression changes of the genes containing three or more CUG repeat tracts in tibialis anterior (TA) of ARC-S and PBS-injected HSA_LR_ mice (n=3). **c,** GO analysis of differentially expressed genes between iAAV9.6-ARC-S and PBS treated HSA_LR_ mice. *P* values were calculated using modified Fisher’s exact test; −log_10_-adjusted *P* values are indicated. **d,** Hierarchical clustered heatmap shows normalized gene expression levels of muscle-related genes. Identification of differentially expressed genes (DEGs) in skeletal muscle (HSA_LR_ *vs.* WT) (top). Expression patterns of identified DEGs across experimental groups in HSA_LR_ mice (n=3) (bottom).

**Extended Data Fig. 10.**
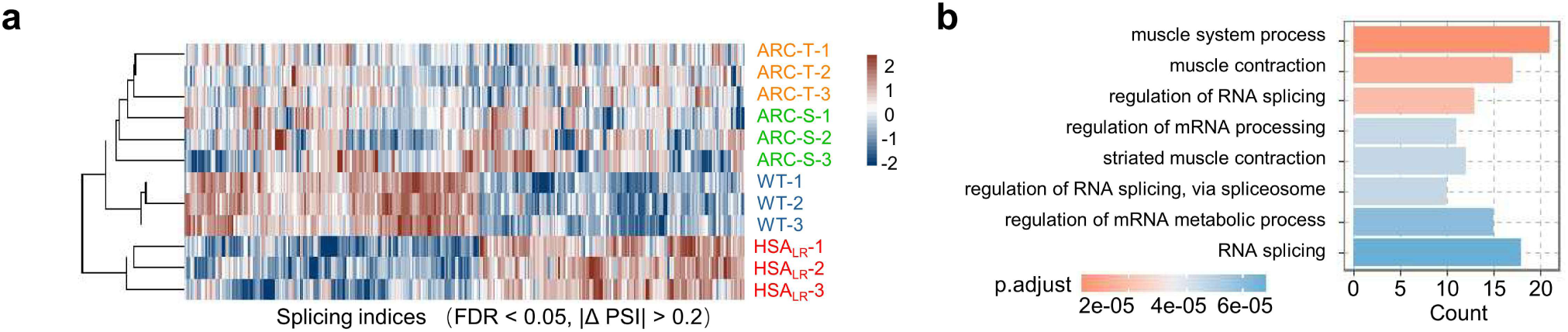
Functional annotation of splicing changes in tibialis anterior of HSA_LR_ mice. **a**, Hierarchical clustered heatmap of exon inclusion indices in AAV-treated tibialis anterior (n=3 for each group). **b,** GO analysis of altered splicing events between iAAV9.6-ARCs and PBS treated HSA_LR_ mice. *P* values were calculated using modified Fisher’s exact test; −log_10_-adjusted *P* values are indicated.

**Extended Data Fig. 11.**
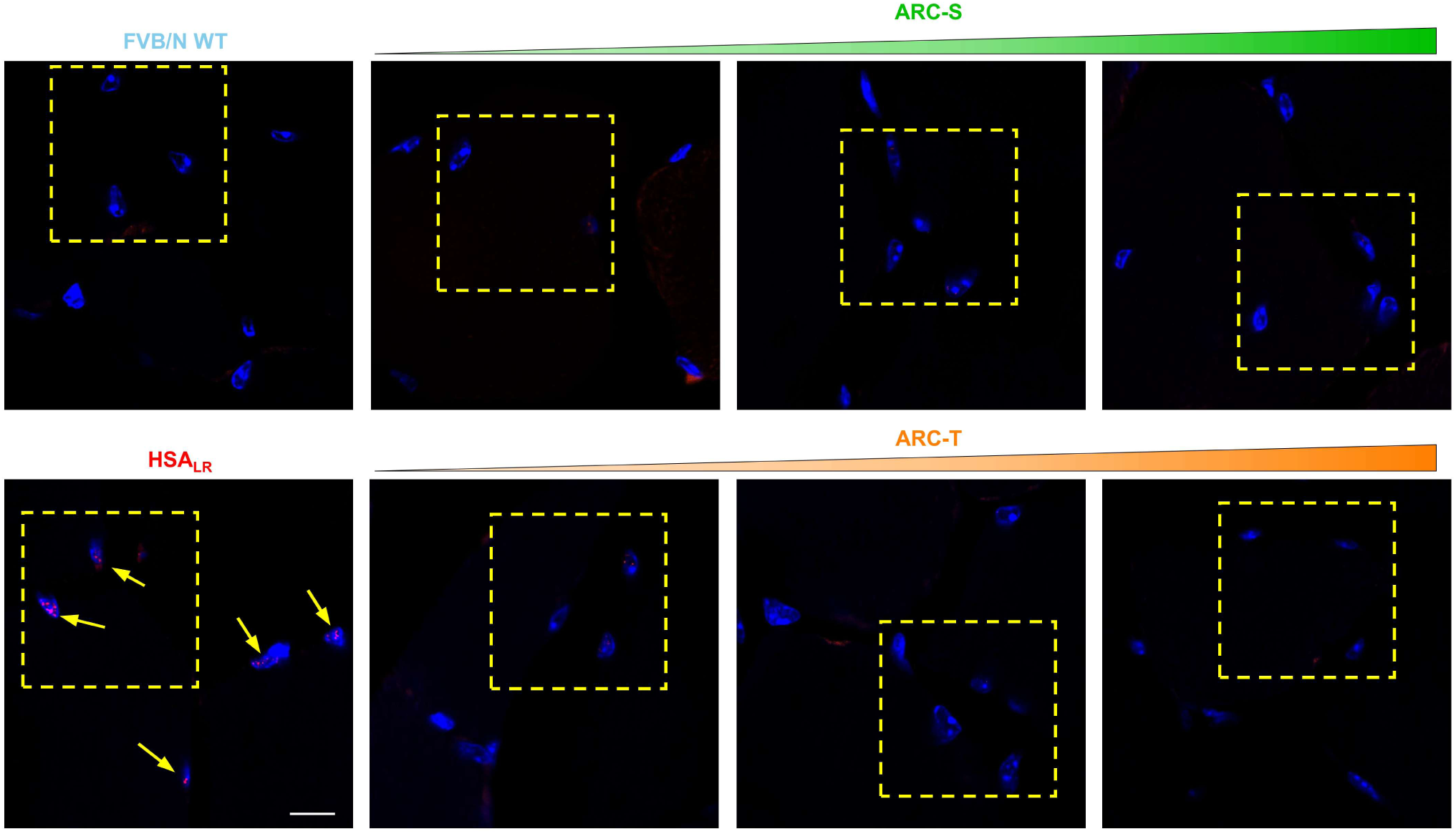
iAAV-9.6-ARCs reduce RNA foci in TA muscle of HSA_LR_ mice. Full-field image of the muscle section. The square area (yellow box) is enlarged and shown in Fig. 6b. RNA-FISH analysis of CUG repeats (red) and cellular nucleus (DAPI, blue) of latitudinal sections of TA muscle treated with ARCs (representative image, n = 3). The arrowheads highlight the presence of CUG RNA foci (yellow). Scale bars, 10 µm.

## Acknowledgements

We thank Ms. Yun Jiang for helps in preparing the paper, and members of Wang Lab and Mao Lab for discussion and comments of this manuscript. This work is supported by National Natural Science Foundation of China (32030064 and 32250013 to Z.W., 32471511 and 31971367 to M.M.), Science and Technology Commission of Shanghai Municipality (24ZR1476500 to M.M.), BMGF-NSFC Joint Project on Vaccine Research and Development (W2412022 and 2024VTJP1001 to Z.W), the National Key Research and Development Program of China (2021YFA1300503 to Z.W) and the Strategic Priority Research Program of Chinese Academy of Sciences (XDB38040100 to Z.W).

## Author Contributions

Conceptualization, M.M. and Z.W.; Methodology, T.W., M.M. and Z.W.; Formal Data Analysis, T.W. and M.M.; Investigation, T.W., M.M., H.H, X.J., C.Y., H.L., W.H.; Writing-Original Draft, T.W., M.M. and Z.W.; Writing-Review & Editing, T.W., M.M. and Z.W.; Funding Acquisition, M.M. and Z.W.

